# Stimulus dependent diversity and stereotypy in the output of an olfactory functional unit

**DOI:** 10.1101/133561

**Authors:** Ezequiel M. Arneodo, Kristina B. Penikis, Neil Rabinowitz, Annika Cichy, Jingji Zhang, Thomas Bozza, Dmitry Rinberg

**Affiliations:** NYU Neuroscience Institute, New York University Langone Medical Center, New York, NY; Center for Neural Science, New York University, New York, NY; Howard Hughes Medical Institute, New York, NY; Department of Neurobiology, Northwestern University, Evanston, IL

## Abstract

Olfactory inputs are organized in an array of parallel functional units (glomeruli), each relaying information from sensory neurons that express a given odorant receptor to a small population of output neurons, mitral/tufted (MT) cells. MT cells have complex temporal responses to odorants, but how these diverse responses relate to stimulus features is not known. We recorded in awake mice responses from “sister” MT cells that receive input from a functionally-characterized, genetically identified glomerulus, corresponding to a specific receptor (M72). Despite receiving similar inputs, sister MT cells exhibited temporally diverse, concentration variant, excitatory and inhibitory responses to most M72 ligands. In contrast, the strongest known ligand for M72 elicited temporally-stereotyped, early excitatory responses in all sister MT cells that persisted across all odor concentrations. Our data demonstrate that information about ligand affinity is encoded in the collective stereotypy or diversity of activity among sister MT cells within a glomerular functional unit in concentration-independent manner.

## INTRODUCTION

Objects in the world are represented by complex patterns of activity in peripheral sensory neurons. Prior to reaching cortical areas, these representations are transformed and reformatted. One of the central challenges in sensory neuroscience is to understand the functional role and computational logic of these transformations in extracting salient information about the environment.

In mammals, the olfactory bulb (OB) is the single interface between primary olfactory sensory neurons (OSNs) and higher brain regions such as piriform cortex. OSNs carry information about odors into the olfactory bulb via a vast array of glomeruli. Each glomerulus is a functional unit, collecting input from OSNs that express a single olfactory receptor gene^1^ and that share similar response properties^2^. Each glomerulus provides exclusive excitatory input to a set of 10-20 mitral/tufted (MT) cells, which project to higher brain areas^3^. The output of a given MT cell depends not only on the response of the glomerulus providing its input, but also on the activity of the complex network of inhibitory interneurons within which it is embedded^3^.

It is still not understood how odor information is represented in MT cells. As an odor is inhaled, a unique subset of glomeruli is activated, resulting in a spatiotemporal pattern that evolves over the course of the respiration cycle^4,5^. Once this input reaches the MT layer, however, there is substantial heterogeneity among cellular responses. The population of MT cells responds to a given odor with various combinations of temporally patterned excitation and inhibition^6,7^. Recent observations from anaesthetized animals suggest that MT cells that are connected to the same glomerulus (sister MT cells) respond to odors with diverse signs of deviation from baseline and timing^8-10^. However, it is not clear how the complexity and diversity of MT responses relate to specific attributes of the odor stimulus. What determines whether sister MT cells show uniform or divergent responses to a given odorant? Are these response properties stable under natural variation in the odor signal, such as changes to odor concentration? Given that sister MT cells do not always behave in a unified way, what information can this subpopulation of cells convey about an odor?

Here, we provide an answer to these questions by functionally dissecting a glomerular functional unit in awake mice. Using a combination of mouse genetics, electrophysiology and imaging, we define the functional properties of inputs to a genetically-tagged glomerulus, and then use optogenetics to identify MT cells that get input from this glomerulus. We observe, for the first time, stimulus dependent diversity and uniformity in sister MT cell responses in awake animals, and we find that relative ligand affinity for an odorant receptor is a major determinant of whether the MT cells responded in a uniform manner manner and whether a cell’s response is consistent across concentrations. Our results directly link a fundamental stimulus property with a robust, concentration-invariant response feature, and suggest a novel way of looking at olfactory coding.

## RESULTS

To study how a single channel in the olfactory bulb processes stimulus information, we characterized the inputs and outputs of the mouse M72 glomerulus. First, to characterize the input, we measured the responses of genetically identified M72-expressing OSNs (M72-OSNs) to a defined odorant set of M72 ligands in a semi-intact preparation of the olfactory epithelium^11^. The dendritic knobs of fluorescently labeled OSNs from M72-GFP mice^12^ were targeted for recording via perforated patch (Fig. 1a). The relative sensitivities of M72-OSNs to each ligand covered a large range of receptor sensitivities: EC_50_ values (concentration at half-maximal response) of the seven odorants spanned three orders of magnitude, from 0.03 to 36 μM (Table 1, Fig. 1b, c). In all figures, we present odors rank-ordered by the M72-OSN sensitivity, from least sensitive (high EC_50_) on the left, to most sensitive (low EC_50_) on the right.

**Table 1.**
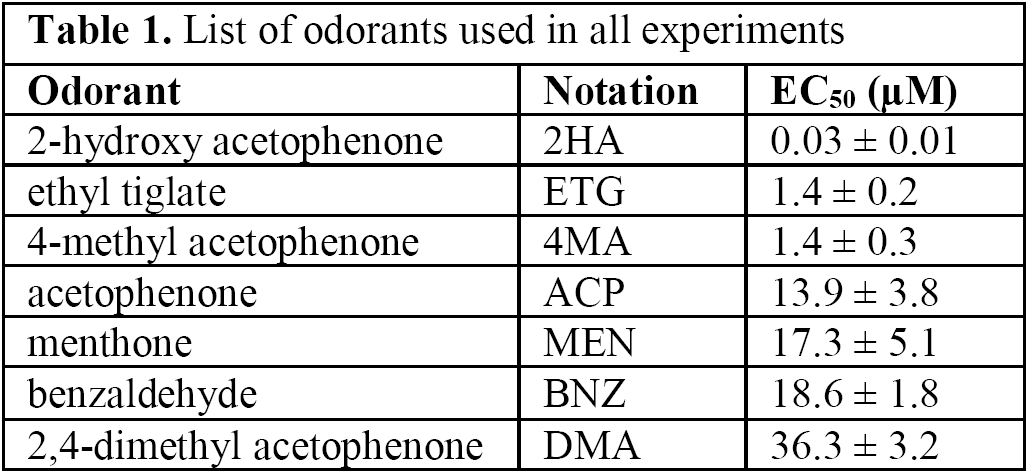
List of odorants used in all experiments

**Figure 1.**
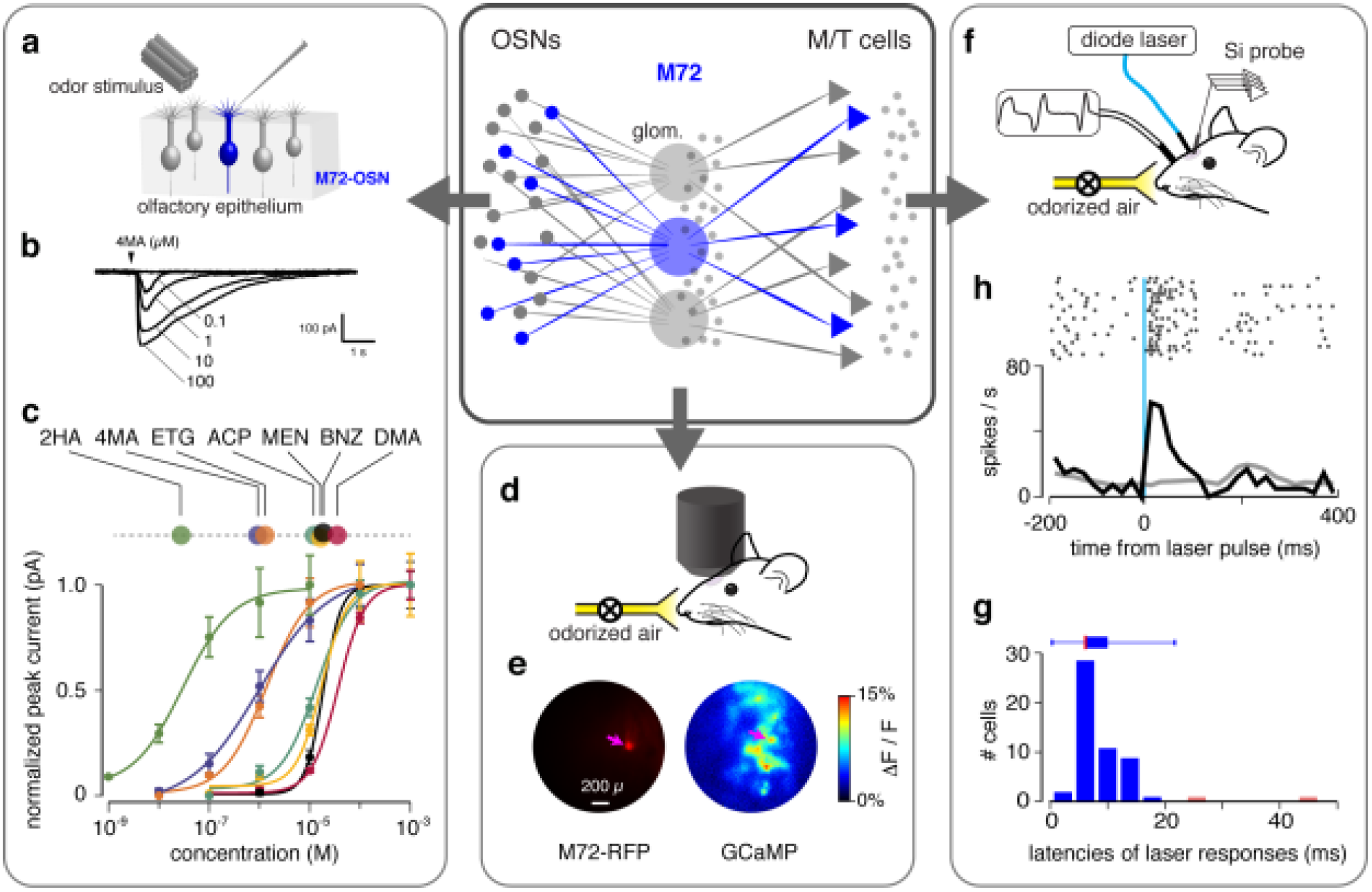
Characterizing information in a single channel of the mouse olfactory bulb**. Central Insert:** Schematic of the olfactory bulb network. Axons from olfactory sensory neurons (OSNs) expressing the same olfactory receptor gene converge to form glomeruli, and each glomerulus provides the sole excitatory input to a few MT cells. Odor signals are subject to significant modification by a network of inhibitory neurons (small gray dots). **(a)** Experimental setup for characterizing OSN responses to odor. Patch clamp recordings are made from dendrites of fluorescently labeled OSNs expressing the M72 odorant receptor. **(b)** Example traces of OSN odor responses. **(c)** Normalized dose-response curves for 7 M72 ligands fitted to the Hill equation (n=5-7 OSNs per odorant; mean ± SEM). EC_50_ values are indicated in linear plot above. Odorant abbreviations and EC_50_ values are given in Table 1. **(d)** Experimental setup for imaging. An awake, headfixed mouse (OMP-GCaMP + M72-RFP) with implanted window above the OB is positioned under the microscope. **(e)** *Left* - an image of RFP M72 glomerulus. *Right –* an example image of glomerulus Ca response to an odor (2HA). M72 glomerulus here and further is marked by magenta arrow. **(f)** Experimental setup for *in vivo* recording of odor responses from MT cells connected to the M72 glomerulus. A head-fixed mouse is positioned in front of the odor port. The sniff signal is recorded by a pressure sensor via a cannula implanted in the nasal cavity. Brief pulses of blue light are delivered to the ChR2-expressing M72 glomerulus through an optical fiber positioned above the glomerulus. MT cell responses are recorded with a multi-shank Si-probe inserted nearby. **(h)** Example of MT cell excitation following laser stimulation of the M72 glomerulus. Raster plot (upper panel) and PSTH (lower panel) around the onset of a 1 ms laser pulse showing the stimulus response (black line) and the baseline activity (grey line). **(g)** Distribution of light response latencies to a light pulse stimulation of 1 ms duration and 5-10 mW power; light responsive cells with latencies longer than 20 ms (colored light red in the histogram) were excluded from the analysis.

Second, to confirm that the M72 ligands would drive MT cells in vivo, we imaged presynaptic OSN activity in identified M72 glomeruli in awake mice (Fig.1d,e). We used a strain of mice in which all the OSNs express the calcium activity indicator GCaMP3, and in which M72-OSNs also express the red fluorescent protein (RFP). This allowed us to assess the level of activation of M72 (and surrounding) glomeruli for each of the odorant stimuli and concentrations used to record MT cell activity.

Third, to characterize the output of the M72 glomerulus, we measured responses of M72-MT cells to the same odorants. To do so, we developed a novel method to identify these cells in awake, freely-breathing animals (Fig. 1f). We used a strain of mice in which M72-OSNs express a channel-rhodopsin2-YFP fusion protein (ChR2-YFP) and are therefore light-sensitive^13^. We periodically stimulated the M72 glomerulus with a 473 nm light pulse while recording extracellular activity in the olfactory bulb (Fig. 1f). Those cells whose firing rate increased shortly after light stimulation were considered putative M72-MT cells (Fig. 1h). The distribution of light-evoked response latencies has its mode and median at 6 ms (Fig. 1g, see Methods); we exclude cells with latencies slower than 20 ms as likely being more than one synapse from the M72 glomerulus. In total, we recorded N = 53 M72-MT cells and N = 312 generic (i.e., non-M72) MT cells.

### Functional characterization of MT cells

MT cell activity is strongly influenced by the temporal patterns of respiration^6,14^ and the duration of odor exposure. In freely-breathing, head-fixed mice, there is considerable variability in sniff frequency and duration (Fig. 2a,b,c). Such variability causes peri-stimulus time histograms (PSTH) of MT cell odor responses to be temporally smeared (Fig. 2c)^6^, and makes it difficult to compare MT responses between different mice with different sniff patterns. Here, we focused our analyses on slower sniffs— those with an inhalation duration >100 ms (Fig. 2c)—because they comprised 75% of all sniffs across all mice, while the rarer, fast sniffs seemed to mark a distinct behavioral state (Fig. S2). Additionally, to avoid adaptation effects, we restricted our analyses to the first sniff cycle after odor onset.

**Figure 2.**
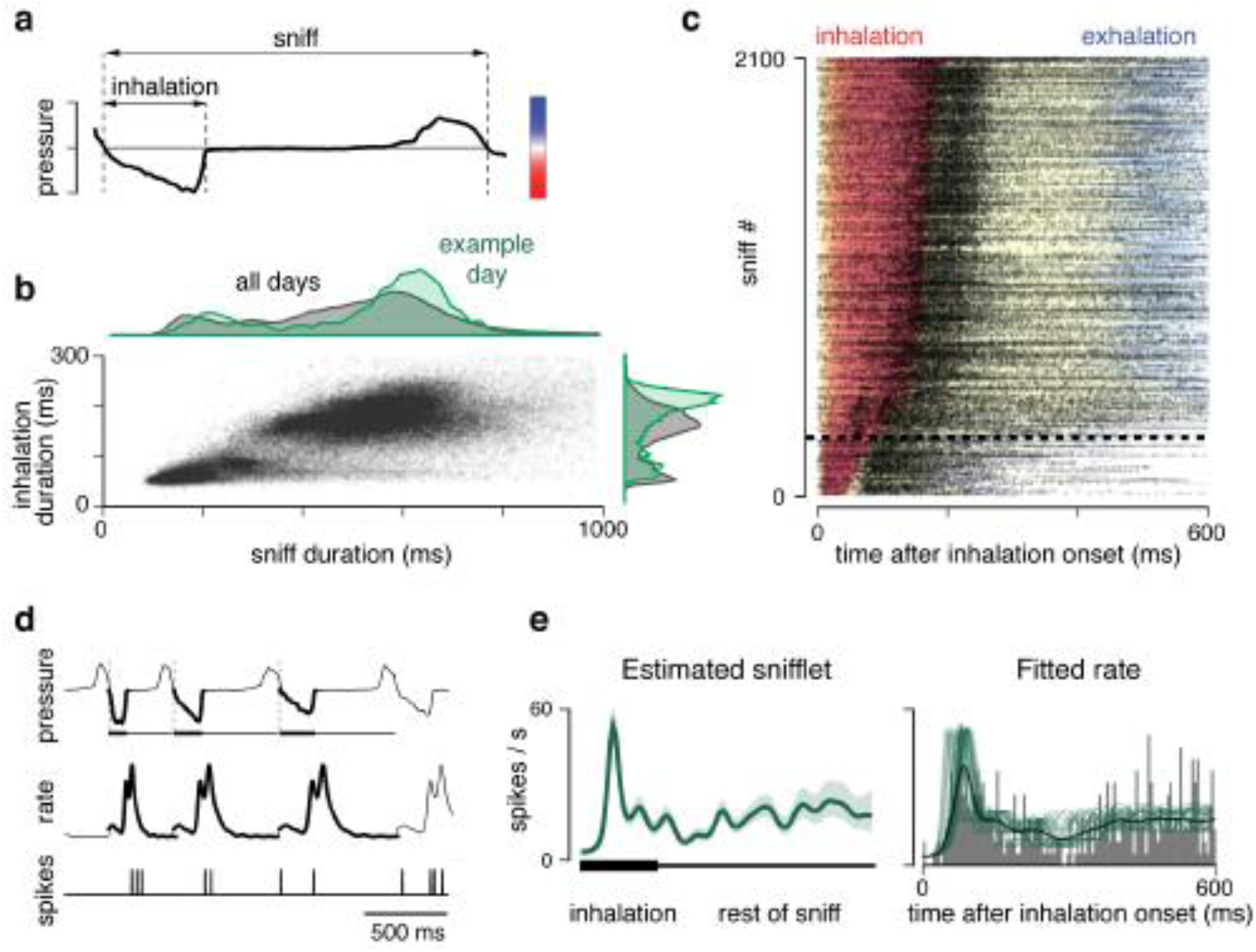
MT cell activity depends on sniff dynamics. **(a)** A pressure signal showing one complete sniff cycle, recorded from the mouse nasal cavity. **(b)** Scatter plot of all inhalation and sniff durations, collated across mice. Top: marginal histogram of sniff durations, across all sessions and mice (black), and an example of one session (green). Right: marginal histogram of inhalation durations. **(c)** Spiking of MT cells depends on inhalation duration. Black dots: raster of spike times of an example M72 cell across 2100 inhalations, during baseline (no odor) condition. Responses are aligned to the onset of inhalation, and are sorted (rows) by inhalation duration. Colored background shows sniff phase: red = inhalation, blue = exhalation. The color map is positioned in panel **(a**), aligned with the pressure axis. Horizontal dashed line demarcates the rarer fast inhalations (below) from the more common slower inhalations (above). **(d)** Snifflet model of MT cell responses. Top: three successive sniffs of short, medium, and longer inhalation duration. Inhalation periods shown as thick regions. Middle: a model fit of the sniff-induced firing rate of the MT cell following a particular temporal pattern, denote as a snifflet. Bottom: the observed spikes. The time courses of the snifflets are the free parameters of the model; these are fitted to each cell, for each stimulus condition, given the observed spikes. **(e)** Snifflet fit to a M72 cell’s response to a single odor. Left: estimated snifflet for this cell/odor. Shaded region: +/-1 SEM. Here and further: the time axis for a snifflet is shown as a thick bar corresponding to the normalized duration of inhalation, and thin bar is the rest of the normalized sniff. Right: grey bars show the trial-averaged PSTH; thick black line shows average firing rate across inhalations; thin green lines show the fitted firing rates on each trial, given the dilations induced by different inhalations.

To further account for variability due to sniff dynamics, we developed a statistical model for the responses of MT cells that factored in both the dependency on the stimulus and on the pattern of sniffing. We modeled the spiking response of an MT cell as arising due to an odor-dependent firing rate pattern, a “snifflet”, that gets temporally dilated as a function of the duration of each sniff (Fig. 2d). The model fits are best when the temporal dilation is a function of the inhalation duration (Fig. S3). To characterize how each cell responds to each odor, we estimate the corresponding snifflet from the observed spiking data (Fig. 2e), which we accomplish using fast Bayesian methods^15,16^. This snifflet representation factors out the variability in sniff duration, allowing us to compare activity across cells and across mice.

From this point forward, we characterize the odor-evoked MT cell responses by comparing the corresponding snifflets. Our results, however, do not depend on the specifics of these modeling decisions: when we used the snifflet model without temporal dilation, the results are qualitatively identical (Fig. S4). Likewise, analyzing MT responses during fast sniffs yields the same general trends, but the lower number of events precluded a rigorous analysis (Fig. S5).

### Diversity and stereotypy of MT cell responses

Despite receiving input from OSNs with defined response properties, we observed a striking degree of response diversity across M72-MT cells (columns of Fig. 3). This diversity was observed for most M72 ligands, presented at the same approximate concentration, 0.075±0.01 μM. Interestingly, responses to one particular ligand, 2-hydroxyacetophenone (2HA), were less variable across the M72- MT cell population (right column of Fig. 3). Almost every cell responded to this odor with a short-latency increase in firing rate. After this robust, reproducible burst, the responses diverged and exhibited considerable variability. We note that 2HA is the strongest ligand yet identified for M72, with an EC_50_ that is two orders of magnitude lower than that of any other identified M72 ligand^11^ (Fig. 1c, Table 1).

The differences in M72-MT behavior for 2HA and other ligands cannot be attributed simply to different levels of parent glomerulus activation, since 2HA and other odorants (like DMA, ACP, 4MA) evoked similar amplitude responses in the M72 glomerulus (Fig 4a). Thus direct feedforward activation of M72-MT cells via their parent glomerulus does not alone determine whether their responses are diverse or stereotyped.

**Figure 3.**
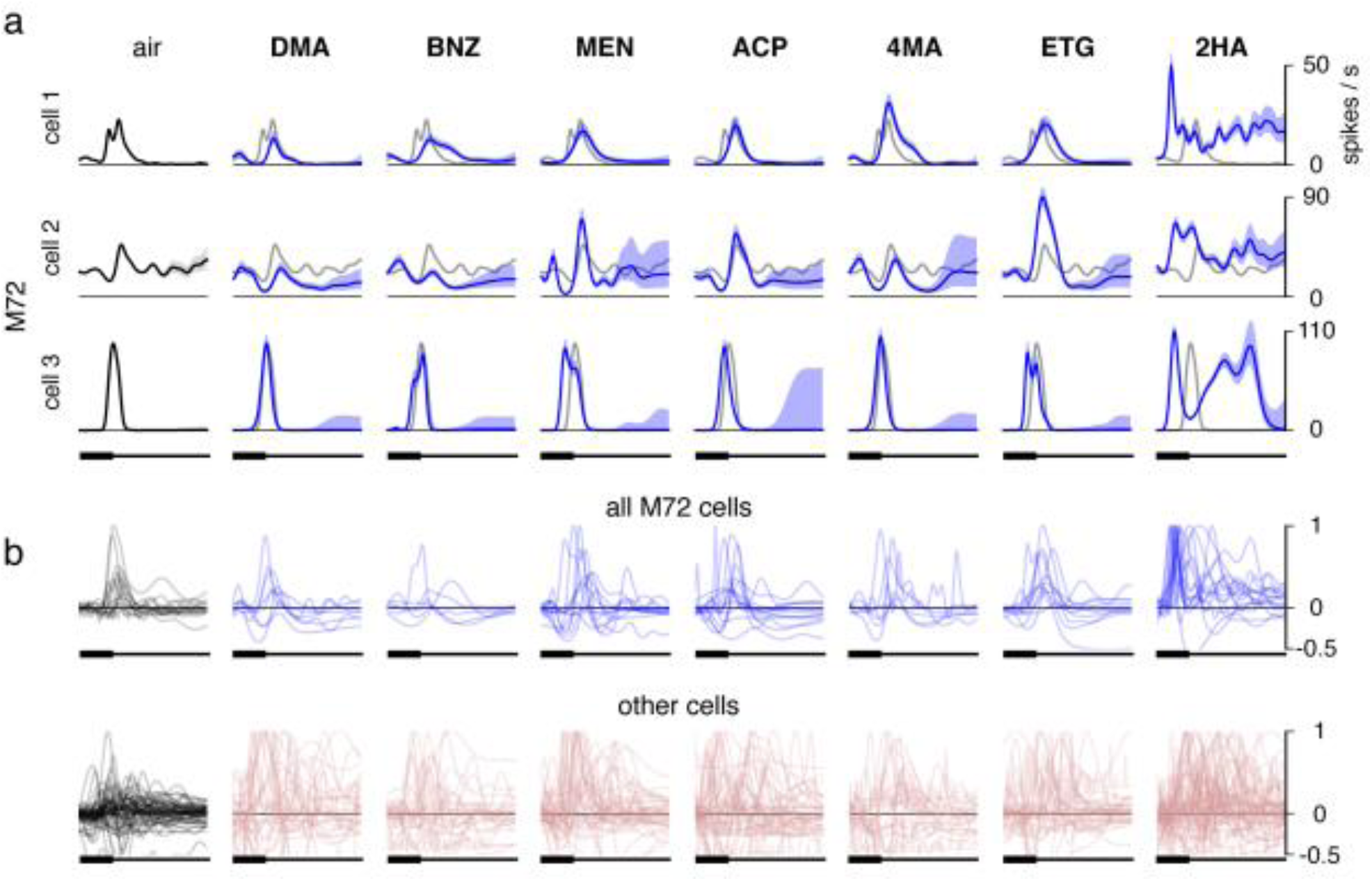
MT responses to odor presentation. **(a)** Example M72-MT cells. Each panel shows the snifflet estimated for each cell and each odor (in blue, sorted by M72 OSN affinity), based on the first sniff after odor onset. First column: baseline, i.e. no odor. Baseline snifflets are estimated from activity three seconds prior to each odor onset. Baseline snifflets are shown in grey in the odor panels for comparison. **(b)** First row: Overlaid snifflets from all recorded M72-MT cells. Snifflets have been normalized on a cell-by-cell basis (so that a cell’s mean snifflet value across odors at inhalation onset is zero, and its maximum peak across odors is unitary). Snifflets are omitted if the cell’s responses were best described as constant over the duration of the sniff. For all odors except 2HA, there is substantial variability in how these cells respond. Second row: the same but for non-M72-MT cells.

To quantify the diversity across cell responses, we constructed several metrics (Fig. 4b-d). First, for each odor, we computed the mean response of the ensemble of MT cells. We normalized each cell’s set of snifflets by a snifflet with largest amplitude and then averaged these across cells. The mean responses of M72-MT cells to most odorants were barely distinguishable from their respective mean responses in the baseline (no odor) condition (Fig. 4b). In contrast, the excitatory response to the strongest ligand (2HA) was still present within the mean activity.

Second, we compared the polarities of the response of each cell to each odor. For each cell and odor, we labeled the response as excitatory or inhibitory, based on the sign of the first significant (3σ) deviation of the cell’s odor-evoked snifflet from its baseline (i.e. no odor) snifflet (Fig. 4c, left). For all odors except 2HA, there was considerable diversity amongst M72-MT cells in first response polarity, with roughly one third of cells having an excitatory response, one third having an inhibitory response, and the remainder being unresponsive to the odor (i.e. no significant deviation from baseline). These distributions were indistinguishable from those found amongst the generic MT cell population (Fig. 4c, right; Pearson’s chi-squared tests). Again, the exception was 2HA, for which almost every M72 cell’s first significant response was excitatory.

**Figure 4.**
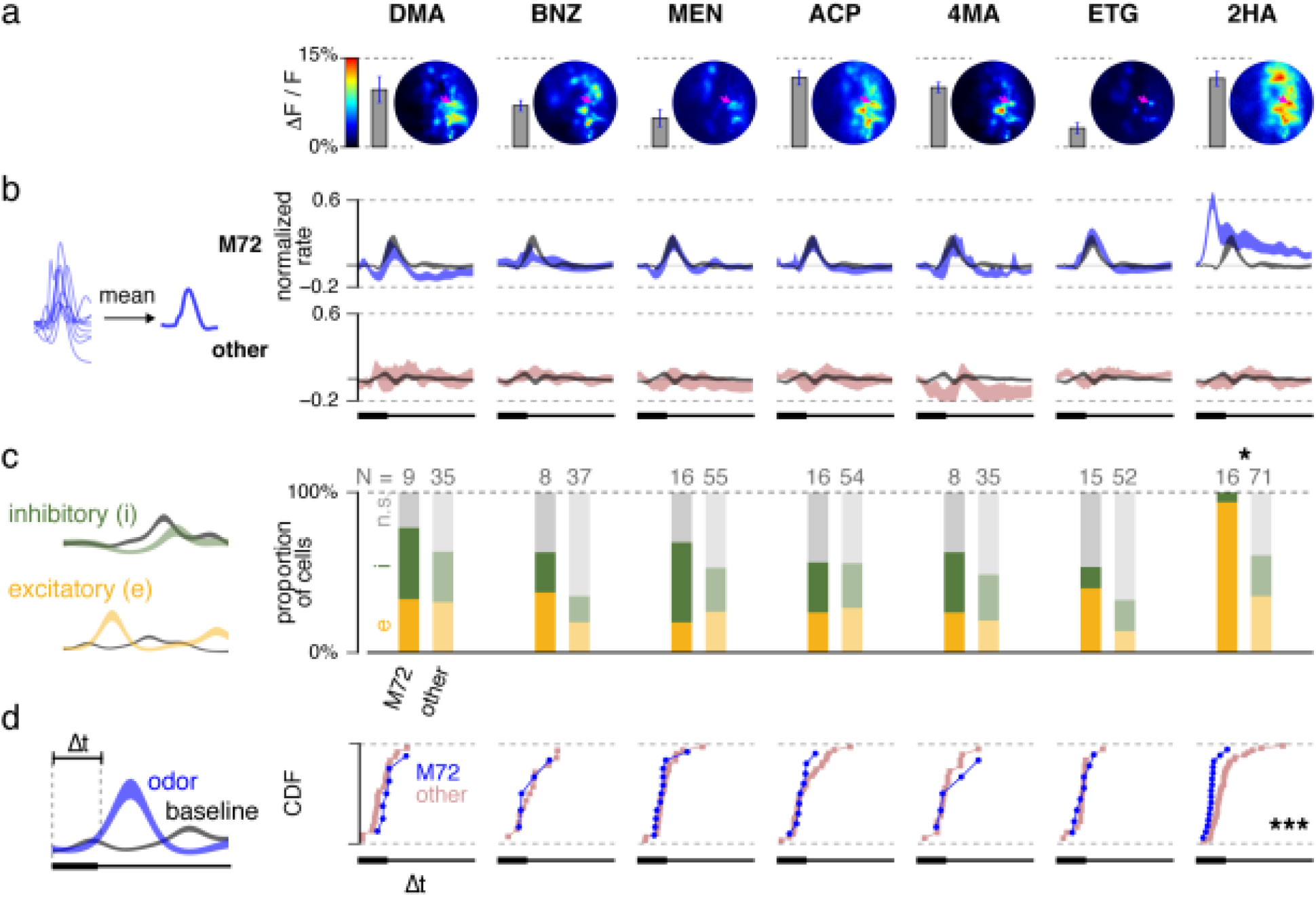
Response variability/stereotypy across the M72-MT population. (a) Map of Ca activity of OSN terminal in the vicinity of M72 glomerulus (pink arrow) for all odors. Vertical bars indicate the level of M72 glomerulus activation for each odor. **(b)** Mean normalized snifflets for each odor (as in Fig 3b), averaged across cells. Left (for all following): schematic of measurement. Right: mean snifflets. M72-MT cells in blue; generic MT cells in pink; mean baseline snifflets in grey. Shaded region denotes ±1 S.E.M. **(c)** Polarity of the first significant response. Right: histograms of first response polarity (yellow: excitatory; green: inhibitory; grey: no significant rate change) for M72-MT cells (bold) and generic MT cells (desaturated). Statistical tests: Pearson’s chi-squared, comparing number of excitatory/inhibitory responses for M72 and generic cells (here and further: * denoting p < 0.05, ** for p < 0.01, and *** for p < 0.001). **(d)** Latency to first significant response. Significance is measured as a 3**s** difference between the odor-evoked snifflet and the baseline snifflet. Right: cumulative distribution functions (CDF) of latencies amongst the M72 (blue) and generic (pink) MT populations. Cells for which odor-evoked activity did not deviate from baseline are omitted. For all odors but 2HA (last column), there is no significant difference between the latency distributions of the M72-MT population and of the generic MT population (Kolmogorov-Smirnoff 2-sample tests).

Third, we compared the onset latencies of odor responses. We computed these as the time of first significant deviation of a cell’s odor-evoked snifflet from its baseline snifflet (Fig. 4d, left). For all odors but 2HA, the distributions of latencies for M72-MT cells were indistinguishable from those seen amongst the generic MT cell population (Fig. 4d, right; Kolmogorov-Smirnov tests). For 2HA, response latencies amongst M72-MT cells were consistently short (Fig. 4d).

In summary, although the M72-MT cells receive common input from sensory neurons in a glomerulus, their responses to any given odor are typically as diverse as the rest of the MT population. The exception to this pattern is a high affinity M72 ligand, 2HA, to which M72-MT cells respond with an initially stereotyped temporal profile, characterized by a strong, short-latency, excitatory transient.

### Diversity and stereotypy of the M72-MT population response is not a concentration effect

The experiment above reveals two different response modes for the M72-MT population: cells can either respond with similar temporal profiles (as we see for 2HA), or with a diverse range of temporal profiles (as we see for all other odors). But which feature of odor stimuli determines the population response mode? Is it the *identity* of a stimulus (i.e. ligand strength) or the *effective concentration* of a stimulus?

To address this question, we selected two odors – menthone (MEN), a weaker ligand, and 2HA, the strongest ligand – and presented them at concentrations spanning 2 orders of magnitude (N = 14-16 M72, N = 107-167 generic MT cells; not every cell tested on every odor/concentration condition). We decreased the concentration of 2HA 10-fold (C_-1_) and 100-fold (C_-2_), and both increased and decreased the concentration of MEN 10-fold (C_+1_ and C_-1_, respectively). This allowed us to decouple the odor-specific and concentration-specific factors that might determine the population response mode.

As the concentration of MEN changed, the level of M72 glomerulus activation varied from almost no response at lowest concentration to a near saturating response at the highest concentration (Fig. 5a). Despite this significant change in glomerular activation, the M72-MT responses remained as diverse as the generic MT responses (Fig. 5b-d, left). In contrast, M72-MT responses to 2HA remained stereotyped at all concentrations (Fig. 5b-d, right). Thus, the diversity of M72-MT cell responses is dependent on odor identity, and not concentration.

**Figure 5.**
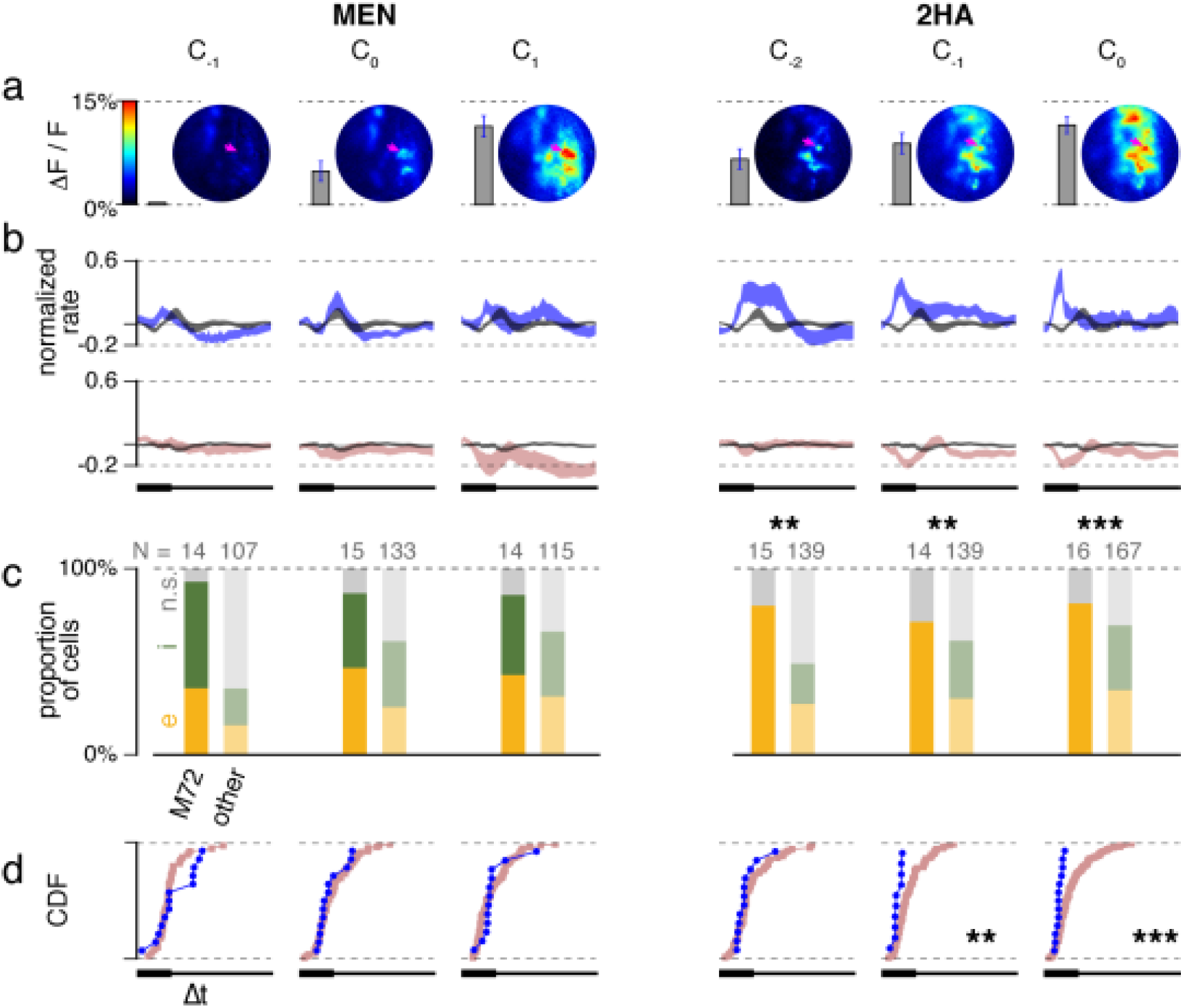
Robustness of results across odor concentrations, spanning 3 orders of magnitude. Panels as in Figure 4. Left three columns show increasing concentrations of MEN, a weak ligand for the M72-OSNs. Right three columns show increasing concentrations of 2HA, a strong ligand for the M72-OSNs. C_0_ denotes concentrations equal to that measured in Figure 4; C_1_ denotes a 10x increase in odor concentration. (Note: due to the differences in experimental setups for imaging experiments the concentrations for MEN were 1.8x lower than the correspondent concentrations in electrophysiological experiments and 2x higher for 2HA.)

### Stereotyped responses to the strongest ligand are robust to concentration changes

Thus far, we have shown that the M72-MT cell population responds in a stereotyped way to a strong ligand, but with considerable diversity to other ligands. This raises the question whether *individual MT cells* respond in a different manner to these two classes of stimuli.

Analyzing individual cells from the second dataset above, we found that changing the concentration of a single odorant could affect single MT cell responses in different ways. As shown in Fig. 6a, the responses of two M72-MT cells to 2HA were consistent at different concentrations: these cells displayed an early excitatory transient at all three concentrations, with the onset latency decreasing as concentration increased, similar to recent reports^17^. Conversely, responses of the same cells to MEN significantly changed with concentration (Fig. 6a, left): increasing the concentration of MEN often diminished, vanished or reversed an excitatory response observed at a lower concentration.

**Figure 6:**
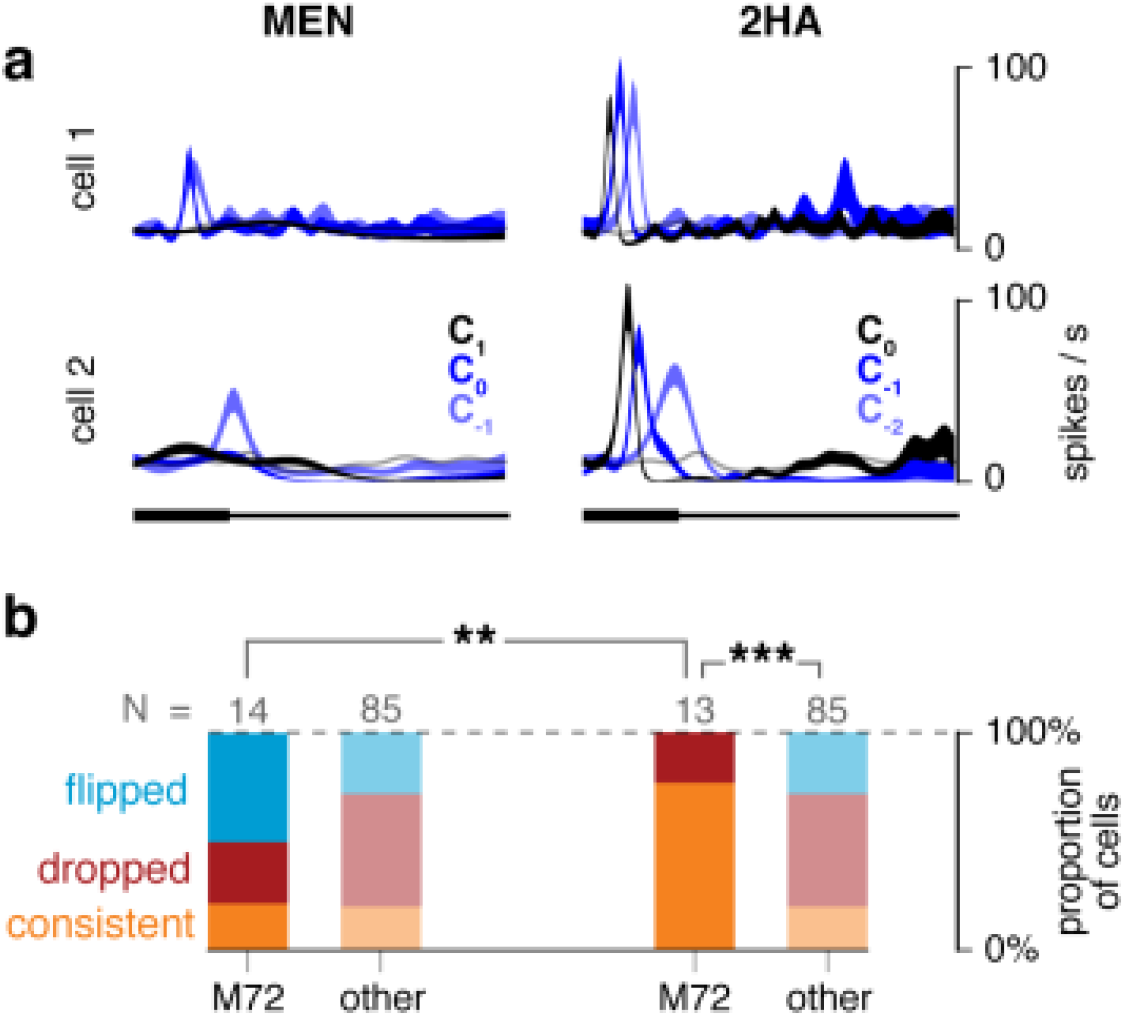
M72-MT cells provide a concentration-tolerant code for 2HA, but not MEN. **(a)** Odor responses of two example M72-MT cells to three concentrations of MEN (weak ligand) and 2HA (strong ligand). Shaded areas denote ±1 S.E.M.; light grey trace shows baseline response. Both example cells respond to increasing concentrations of 2HA with an initial excitatory response, and a latency that decreases with concentration. Neither cell responds to the highest concentration of MEN. **(b**) Distribution of cells by category of concentration dependency for MEN (left) and 2HA (right). We define three categories: *consistent* (orange) – the response polarity is the same for all three concentrations; *dropped* (red) – there is at least one concentration at which the response did not significantly differ from baseline; and *flipped* (blue) – the response polarity is different at different concentrations. Cells for which the response was not significantly different from baseline in all three concentrations are omitted. Pairwise comparisons: chi-squared tests.

To quantify these observations, we categorized the concentration dependency of each cell’s response to each odor, based on the variability of its first significant deviation from the baseline response. We assigned the label *consistent* if the cell’s first significant response to the odor was always excitatory or always inhibitory, across concentrations; *dropped,* if there was a response for one or two of the concentrations, but no significant response for the other(s); or *flipped,* if responses with distinct polarities were observed at different concentrations. We found that the majority of M72-MT cells had consistent initial responses across concentrations of 2HA, while the distribution of response categories was mixed for MEN and for the generic MT population (Fig. 6b).

Thus, the strong ligand not only elicits stereotypic responses across M72-MT cells, but also evokes temporal response patterns within cells that are robust to changes in concentration. These observations do not appear to hold for other ligands of M72, nor amongst cells of the general MT population. These observations identify MT response features that are concentration-invariant.

## DISCUSSION

Glomeruli are considered functional units in early olfactory processing. Here, we have studied for the first time the odor-evoked responses in MT cells that receive input from a genetically-identified glomerulus in awake, freely-breathing animals. We optogenetically identified sister MT cells that receive excitatory drive from the glomerulus of the M72 odorant receptor, recorded their responses to a set of well-characterized M72 ligands, and analyzed the data using novel statistical tools (Figs. 1-2). Despite receiving excitatory drive from functionally-similar sensory neurons, M72-MT cells responded to most odorants with highly diverse temporal patterns that were as heterogeneous as those found in the generic MT population. However, this response diversity was not observed in response to 2HA, which has the highest apparent affinity of all identified M72 receptor agonists (Figs. 3-4). M72-MT responses to 2HA almost always included a stereotyped early excitatory transient, after which response diversity resumed. These odor-specific patterns of response diversity and stereotypy remained unchanged across odor concentration (Figs. 5-6), suggesting that they do not depend on how strongly the glomerulus is activated. Our data demonstrate that MT cells within a specific olfactory functional unit encode a strong ligand in a markedly different way than weaker ligands.

Previous studies^6,7^ have demonstrated that responses amongst randomly selected MT cells in awake mice are diverse in their polarity and timing across the sniff cycle. Our results suggest that this diversity cannot be attributed solely to different sources of glomerular feedforward input, as we observed that MT cells sharing a common glomerulus show similar response diversity to most odorants.

### Sources of response variability

There are multiple potential sources for MT cell response diversity: 1) variability across animals; 2) variation in intrinsic biophysical properties across cells^18,19^, including differences between mitral and tufted cells^20^; 3) non-homogeneity of excitatory synaptic connections within a glomerulus; and 4) heterogeneity of inhibitory network connectivity. While the first three factors may play some role in the observed diversity of the responses, they cannot easily explain the fact that this diversity vanishes with a strong ligand, nor can they explain that a strong ligand (but not a weak ligand) evokes stereotypical responses independent of concentration.

A significant source of between-cell response variability comes from the rich inhibitory network, which includes granule cells, periglomerular and other inhibitory cells in the olfactory bulb. The connections from granule cells to MT cells are sparse and heterogeneously distributed^3,21,22^, as are the connections from the periglomerular inhibitory network^23,24^. Functionally, too, the activity of each MT cell appears to be influenced by a handful of sparsely-distributed glomeruli^25^. Diversity across M72-MT cell responses to a single odor could therefore result from each cell having different connectivity within the lateral network.

### The origin of response diversity and stereotypy in sister MT cells

We propose that the relative timing of activity entering the olfactory bulb might be responsible for the observed diversity and stereotypy of responses in sister MT cells. Increasing odorant concentrations evokes shorter latency responses in MT cells^17,26,27^. Consequently, when an odor is presented, OSNs whose receptors have highest affinity to the odorant may be excited first (Fig. 7a). One plausible mechanism for this is a gradual rise in odorant concentration during inhalation, over tens to hundreds of milliseconds, resulting in different OSN types reaching threshold at different times^28,29^, or it could be due to processes of signal integration within the receptor neurons themselves^30^. In turn, a sequence of OSN activation latencies will roughly translate into a sequence of glomeruli conferring excitatory drive to downstream MT cells. As this process begins, the glomeruli and post-synaptic MT cells that are activated early will also propagate signals into the inhibitory networks. This network activity will then feed back into the responses of the population of MT cells in different ways (due to the heterogeneity of connections), thus diversifying subsequent responses (Fig. 7b). For a given odorant, the net effect of these feedforward and recurrent dynamics will thus be different for MT cells in early-and late-responding channels. The initial input experienced by MT cells of early-responding channels will be dominated by feedforward excitation, causing these cells to produce a stereotypical burst of action potentials early in the sniff cycle (Fig. 7c). MT cells associated with late-responding channels, will receive heterogeneous inputs from the inhibitory network coincidentally with the feedforward excitation, resulting in diverse responses within sister cells in the channel.

**Figure 7:**
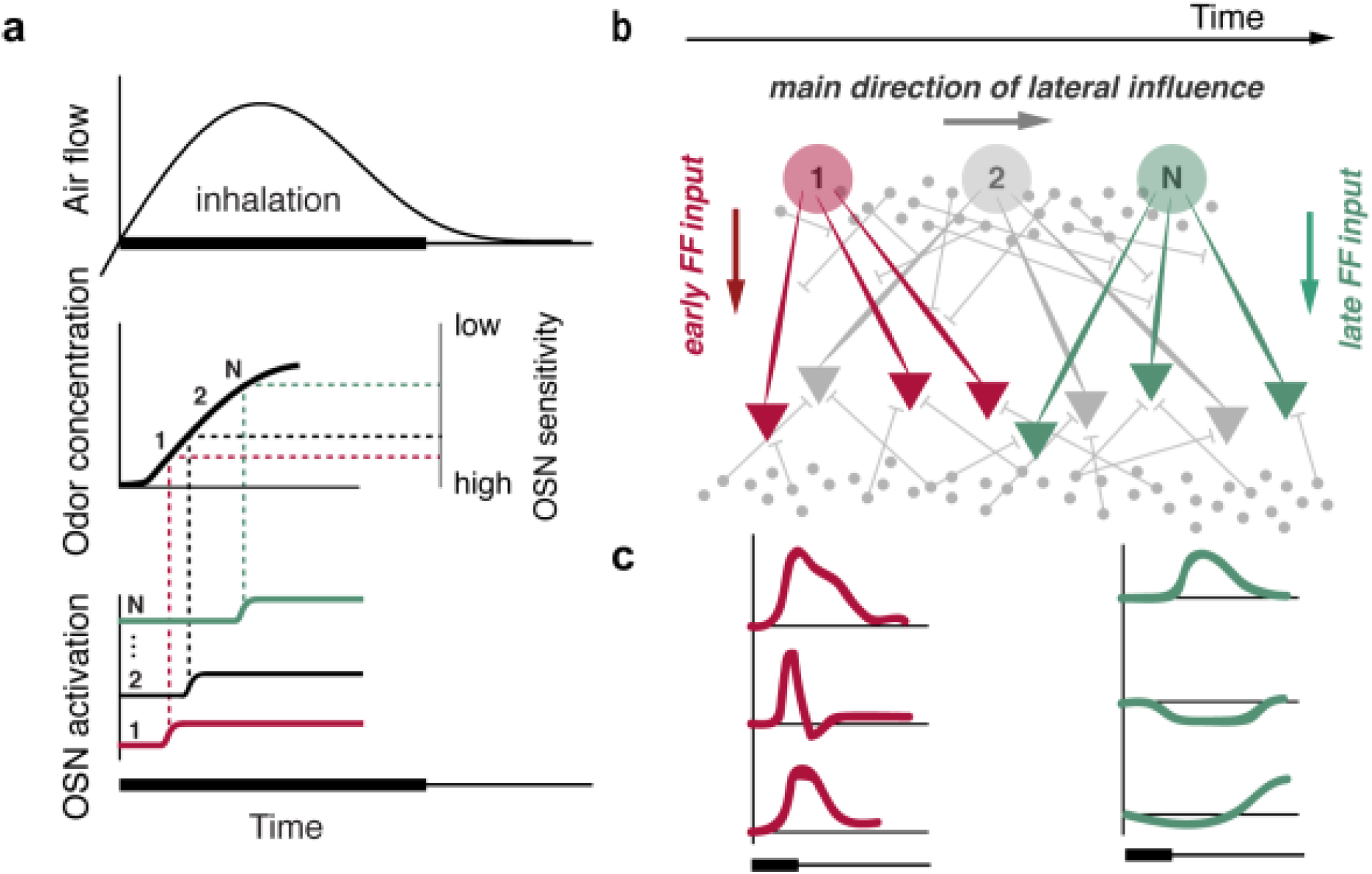
A proposed mechanism responsible for stereotypy/diversity amongst sister MT cell odor responses. **(a)** In the presence of a given odor, the OSN receptors’ relative sensitivities to the odorant determine the relative response latencies of the corresponding glomeruli. As inhalation drives the odor into the nose (top), odor concentration rises gradually (middle left). More sensitive olfactory receptors reach their activation threshold earlier (middle right), and thus respond earlier (bottom). The labels 1 to N denote the temporal sequence of active OSN channels. **(b)** The ensuing flow of activity in the bulb. The schematized olfactory bulb network is shown as in Figure 1: glomeruli (large colored circles), MT cells (colored triangles), and inhibitory neurons (small grey circles; top = periglomerular layer; bottom = granule layer). Thin gray lines represent inhibitory connections. The glomeruli are depicted horizontally in order of their temporal sequence of activation (1 to N), as shown in (a). The MT cells connected to glomerulus 1 (red) receive early feed-forward (FF) excitation, which then propagates through the inhibitory network. MT cells connected to glomerulus N (green) receive both FF excitation and lateral inhibition. **(c)** Typical responses of MT cells connected to early-activated (red) and late-activated (green) glomeruli. Driven by a common excitatory input, MT cells connected to early-activated glomeruli share an initial short-latency excitatory response. MT cells connected to later-activated glomeruli are subject to both excitatory drive and heterogeneous inhibitory influences, and thus show diverse responses.

The results of our experiment are consistent with this hypothesis. Although we do not know the exact timing of M72-OSN activation relative to other channels for each odor, the fact that 2HA is such a strong ligand for the receptor suggests that the M72 channel is one of the first to be activated in response to this odor. In contrast, the weaker sensitivity of M72 receptors to these ligands predicts that the M72-OSNs would be activated relatively late.

Moreover, this model provides a simple explanation for why these results do not change with odor concentration. As odor concentration decreases, OSNs are typically activated later^4^. However, decreasing odor concentration would not change the *relative* timing of glomerular activation within our model. The earliest activated glomerulus at high concentrations would be expected to remain the earliest activated at lower concentrations. The MT cells connected to late-activated glomeruli will receive diverse and concentration dependent inhibitory drive; thus, sister MT cells would not only respond to the odor in a different manner, but also would show variable responses across concentration (Figs. 5-6). To fully test this hypothesis, we would need access to MT cells across many different channels, spanning a range of known affinities to a single odorant—an experiment that is technically very challenging.

There are a few limitations of our approach. First, we use the sensitivity to M72 receptors as a proxy for a ranking order of the M72 glomerulus in the sequence of all glomerulus activations. However, even for the strongest ligand, we do not know if other receptors could have a higher affinity to 2HA than M72. Our observation of stereotypy of 2HA responses and consistency of concentration series are in agreement with an assumption that M72 is one of the first glomeruli activated by 2HA. Second, it is tempting to assume that MT cells should respond earlier for a higher-affinity ligand than for an intermediate or weak ligand. While this relationship is observed in our data (Fig. S6), the absolute latency of response to a given ligand likely depends not only on receptor affinity, but also on the physical and chemical properties of the ligand. For example, a high concentration of a hydrophilic ligand may evoke earlier responses than a weak concentration of a hydrophobic ligand. Thus we avoid drawing conclusions about latency differences across different odors, and focus instead on latency differences for a given odor between cells.

Our observations and model reconcile two inconsistent observations regarding sister MT cells in anesthetized preparations. Tan and colleagues^8^ recorded the total spike counts of MT cells receiving input from the I7 odorant receptor. Their data suggests that the strongest known ligand for that receptor consistently evokes high spike counts in I7 MT cells, but other odorants (not necessarily ligands for I7) typically do not. We suggest there is a parallel with our findings (Fig. S7), implying that these observations could extend to all glomerular channels in the active, functioning bulb. Moreover, we demonstrate the temporal specificity of this effect, namely that the shared MT response to a strong ligand manifests as a temporally-stereotyped early excitatory transient.

Dhawale and colleagues^10^, working in anesthetized mice, found that sister MT cells typically responded to an odor with substantial temporal diversity, yet responded coherently to the common drive provided by optogenetic stimulation of the parent glomerulus. By studying a glomerulus with functionally defined inputs, we show for the first time that the response diversity among sister MT cells vanishes for a ligand that has high affinity for the odorant receptor of the parent glomerulus. We posit a parallel between activation of a glomerulus with a strong ligand and artificial optogenetic stimulation—in both cases, the specific glomerular excitation can proceed uninterrupted to the corresponding MT layer, without being preceded by activity in other channels, producing synchronous activity in the corresponding sister MT cells.

### Temporal, not spatial, contrast enhancement

MT cells in the olfactory bulb are responsible for conveying information about odor stimuli from peripheral receptors to higher brain areas. A long-standing question is how the odor representation is transformed in this process. Several answers to this question have been posited. At the least, the convergent feedforward input to each glomerulus allows MT cells to average the output of many noisy OSNs, thereby improving the signal-to-noise ratio of the representation^31-33^. It has been suggested that inhibitory networks impose inhibition in a center-surround type of configuration, sharpening the representation of odors in the MT layer^34-37^, much like the role of lateral inhibition in the visual^38,39^, auditory^40,41^, and somatosensory pathways^42-44^. Alternatively, these lateral networks might implement a divisive normalization computation that would render the representation concentration-invariant^45,46^.

Our data show that sister MT cells associated with a given glomerulus encode a strong ligand with temporally stereotyped responses, while they encode weaker ligands with marked response diversity. This represents a novel transform that emerges via the olfactory bulb circuitry.

How might this stimulus-dependent stereotypy be incorporated into a broader olfactory code? We imagine a downstream decoder that is particularly sensitive to synchrony between sister MT cells of a common glomerulus. For strong ligands, sister MT cells will all respond with a relatively short-latency excitatory transient, and the decoding neuron will thus fire as well. For weaker ligands, the temporal diversity amongst sister MT cells will fail to drive the decoding neuron. In such a scheme, lateral inhibition preserves information in early-responding channels and scrambles information in late-responding channels. This configuration would act to sharpen and sparsify the odor representation, reducing the dimensionality of the peripheral combinatorial code to one that is dominated by the most sensitive glomerular channels. Moreover, our observation that the temporal stereotypy/diversity of responses is robust to changes in concentration means that the readout representation would also be concentration-tolerant, and could represent odor identity independent of concentration. These are highly desirable features for an olfactory code.

## Acknowledgments

We would like to acknowledge Chris Wilson, Sasha Devore, Dion Khodagholy, Eero Simoncelli for discussions, Kathy Nagel and Alex Koulakov for comments on the manuscript, Admir Resulaj and Gaby Serrano for technical assistance. Thanks to Loren Looger for sharing GCaMP3 plasmids prior to publication. The work was supported by grants from the Whitehall Foundation (D.R.), and NIH/NICDC grants: R01DC014366 and R01DC013797 (D.R.), R01DC013576 (T.B.), and F31DC014903 (K.B.P). E.M.A. was supported by a Pew Latin American Fellowship in Biomedical Sciences, N.R. was supported by the Howard Hughes Medical Institute, A.C. was supported by the German Research Foundation.

## Author contributions

E.M.A., T.B, and D.R. conceived of the study. E.M.A and D.R. built the experimental set-up. E.M.A., K.B.P., N.R., and D.R. shaped the experimental design. T.B. initiated the transgenic approach and generated the gene-targeted mice. E.M.A. and K.B.P performed electrophysiological recording in awake mice. A.C. performed in vivo imaging experiments. J.Z. performed OSN recordings. N.R. developed and performed snifflet analysis. E.M.A., K.B.P., N.R., A.C., T.B. and D.R. wrote the manuscript. T.B. supervised optical imaging and OSN recordings. D.R. supervised the project.

## Competing financial interests

The authors declare no competing financial interests.

## METHODS

### Animals

For electrophysiological experiments, we used adult homozygous *M72–IRES-ChR2-YFP* naïve mice (Strain *Olfr160* ^*tm1.1(COP4*/EYFP)Tboz*^). Data for MT cell *in vivo* odor responses were collected in 17 animals (13 males, 4 females). Animals were 6–10 weeks old at the beginning of experiment and were maintained on a 12-h light/dark cycle (lights on at 8:00 p.m.) in isolated cages in a temperature-and humidity-controlled animal facility. All animal care and experimental procedures were in strict accordance with protocols approved by the New York University Langone Medical Center and Northwestern University Institutional Animal Care and Use Committees.

### OSN electrophysiology

Perforated patch recording were made from the dendritic knobs of fluorescently labeled M72-expressing OSNs as described^1-3^. In short, the olfactory epithelium from neonatal mice was removed and kept in oxygenated ACSF (95% O_2_, 5% CO_2_), containing 124 mM NaCl, 3 mM KCl, 1.3 mM MgSO_4_, 2 mM CaCl_2_, 26 mM NaHCO_3_, 1.25 mM NaHPO_4_ and 15 mM glucose, pH 7.4, 305 mOsm. The epithelium was transferred to a recording chamber at 20–23°C and imaged using an upright fluorescence IR-DIC microscope equipped with a CCD camera and a 40× water-immersion objective. Perforated-patch clamp was performed by including 260 μM amphotericin B in the recording pipette, which was filled with 70 mM KCl, 53 mM KOH, 30 mM methanesulfonic acid, 5 mM EGTA, 10 mM HEPES and 70 mM sucrose, pH 7.2, 310 mOsm. The electrodes had tip resistances ranging from 8– 10 MΩ and liquid junction potentials were corrected in all experiments. Signals were acquired at 10 kHz and low-pass filtered at 2.9 kHz. Odorants were applied via pressure ejection via a multi-barrel pipette placed 20 μm downstream of the cell. Odorants were dissolved in dimethyl sulfoxide (DMSO) and diluted in bath solution to achieve desired concentration.

### Gene targeting

*OMP-GCaMP3*: The coding sequence of GCaMP3^1,4^ was flanked by AscI sites and cloned into a targeting vector for the olfactory marker protein (*omp*) locus^4,5^ so that the coding sequence of OMP is replaced by that of GCaMP3, followed by a self-excising neomycin selection cassette^5,6^. The targeting vector was electroporated into a 129 ES line, and clones were screened for recombination by long range PCR. Chimeras were generated from recombinant clones by aggregation with C57BL/6 embryos.

*M72-RFP (M72-IRES-tauCherry)*: A cassette containing an internal ribosome entry site (IRES), followed by the coding sequence for a fusion of bovine tau and mCherry^6,7^ and a self-excising neomycin selection cassette, was inserted into an AscI site located 3 nucleotides downstream of the M72 coding sequence in an M72 (*olfr160*) targeting vector^7^. The targeting vector was electroporated into a 129 ES line, and clones were screened for recombination by long range PCR. Chimeras were generated from recombinant clones by injection into C57BL/6 blastocysts.

### Olfactory bulb imaging

Awake *in vivo* imaging: Imaging was done in 7-8 week old naïve, male mice that were heterozygous for the OMP-GCaMP3 and homozygous for the M72-RFP allele, and that had been implanted with chronic optical imaging windows and head bars. Mice were first anesthetized with isoflurane (2-3%) in oxygen and administered buprenorphine (0.1 mg/kg) as analgesic; bupivacaine (2 mg/kg) as a local anesthetic at the incision site; and dexamethasone (2 mg/kg) to reduce cerebral edema. The animal was secured in a stereotaxic head holder (Kopf instruments) and the bone overlying the olfactory bulbs was thinned to transparency using a dental drill. Two micro-screws were placed into the skull to structurally support the head-bar. A custom-built titanium head-bar (3mm x 15mm, <1g) was attached to the skull using Vetbond cyanoacrylate glue and cemented in place using dental cement (Dental Cement, Pearson Dental Supply). Black Ortho Jet dental acrylic (Lang Dental Manufacturing) was extended from the head-cap around the thinned bone forming a small chamber. The area overlying the olfactory bulbs was covered with multiple thin layers of prism clear cyanoacrylate glue (Loctite #411) as described ^2-4^.

Following complete recovery from surgery, mice were placed on a water restriction schedule (1 ml/day). After seven to ten days of water restriction, mice were slowly habituated to the imaging setup where they were trained to lick for a water reward. During imaging sessions, mice were positioned on a custom-built wheel and secured with the head-bar in a custom-built holder.

Light excitation was provided using a 200 W metal-halide lamp (Prior Scientific) filtered through standard filters sets for RFP (49008, Chroma) and GFP (96343, Nikon). Optical signals for GCaMP were recorded using a CCD camera (NeuroCCD SM256; RedShirtImaging) at 25 Hz with a 4x temporal binning. Each recording trial was 16 s consisting of a 6 s prestimulus interval, a 4 s odor pulse and a 6 s poststimulus interval. Only one odorant per day was tested to avoid cross contamination of different odorants. Different concentrations of the same odorant were interleaved with clean air trials to identify potential contamination.

### Response maps

Response maps were obtained by temporally averaging the response signal over a 0.5 s window around the time of maximum response, and subtracting a pre-stimulus response baseline (1.6 s window). For low concentrations, stimuli were presented at least three times and averaged to obtain response maps. Images were processed and analyzed in Neuroplex (RedShirtImaging) and Image J (NIH) software.

Response amplitudes were measured from a region of interest (ROI) drawn around the M72 glomerulus in the RFP image. Only the first trial of each odor concentration was used to obtain the response amplitude to avoid potential adaptation effects.

### Implantation surgery

Mice were anesthetized using isofluorane gas anesthesia. A diamond-shaped bar for head fixation^8^, a reference electrode, and a pressure cannula for sniff recording^9^ were implanted. To implant the sniffing cannula, which was a thin 8.5-mm-long stainless capillary (gauge 23, Small Parts capillary tubing), a small hole was drilled in the nasal bone, into which the cannula was inserted, fixed with glue, and stabilized with dental cement. The reference electrode was implanted in the cerebellum. The mice were given at minimum 5 days after surgery for recovery.

### Setup and odor habituation

After recovery, mice entered a regime of water restriction, with 1 ml administered every day. Five days into this regime, the mice were placed in a head-fixation setup for lick training^8-10^. To reduce stress to the animals and movement artifact during recordings, mice were positioned on a running wheel. Mice could stay still or walk on the wheel if desired. The first few sessions were brief (10–20 min) and served purely to acclimate the animals to head fixation and the running wheel. Mice typically remained mostly quiescent after 1–2 sessions of head fixation, after which lick training sessions began. A lick spout was placed in front of the animal, which delivered a droplet of water every time the animal licked it. Mice typically learned the water-rewarding nature of the head-fix setup within 1-3 sessions. We then removed control of the water delivery from the mouse and started delivering 1 out of 7 odors in pseudo-random sequence, with an average inter-stimulus interval of 8 s and stimulus duration of 1000–4000 ms. A drop of water was delivered to the mouse automatically every 3-5 odor presentations. Animals underwent 3-5 sessions of odor exposure (200-400 trials each) of this type before recordings. This procedure served several purposes: i) it reduced the distress of mice in the setup; ii) it reduced the movement artifact during recordings; and iii) it habituated the animals to the set of odorants used in the experiment, thus eliminating any novelty effects.

### Water delivery

Water delivery was based on gravitational flow controlled by a pinch valve (98302-12, Cole-Parmer) connected via Tygon tubing to a stainless steel cannula (gauge 21, Small Parts capillary tubing), which served as a lick tube. The lick tube was mounted on a micromanipulator and positioned near the mouse’s mouth. The water volume was calibrated to give approximately 2.5 μl per valve opening. Licks were detected by the closing of an electrical circuit through the grounded mouse (the circuit was open until the mouse connected the metal cannula to ground).

### Behavioral and stimulus delivery control

All behavioral events (odor and final valve opening, laser stimulation, water delivery and lick detection) were monitored and controlled by a real-time (1 ms), Arduino platform–based, behavioral controller box, developed at Janelia Farm Research Campus, HHMI. In each trial, the behavioral controller read trial parameters, and sent trial results together with a continuous sniffing signal to a PC running a custom-written Python program, Voyeur (partially developed by Physion Consulting, Cambridge, MA). Voyeur is a trial-based, behavioral experiment control and acquisition software that allows behavioral protocols to compute parameters of trials and send them to embedded real-time hardware systems. The Arduino code and Python application source is available as a GitHub repository (search for Voyeur in GitHub). Every stimulus and behavioral event had an associated trigger signal that was sent to the recording system for precise synchronization with neural activity recordings.

### Sniff recording

To monitor the sniff signal, the implanted sniffing cannula was connected to a pressure sensor through an 8-12 cm long polyethylene tube (801000, A-M Systems). The pressure was transduced with a pressure sensor (24PCEFJ6G, Honeywell) and homemade preamplifier circuit. The signal from the preamplifier was recorded together with electrophysiological data on one of the data acquisition channels. The timing of the pressure signal was calibrated with a hot wire anemometer (mini CTA 5439, Dantec Dynamics, Denmark) as in Shusterman, 2011^9^. The time differences between pressure signal and the flow signal during calibration did not exceed 2-3 ms. The cannula was capped when not in use.

### Light stimulation

Light stimulation was produced via a 100 μm multimodal fiber coupled to a 473-nm diode laser (model FTEC2471-M75YY0, Blue Sky Research). The end of the fiber was cut flat and polished. The light stimulus power at the open end was measured by a power meter (Model, PM100D, Thorlabs), and calibrated to adjust the amplitude of the voltage pulses sent to the laser, to achieve a consistent power output across experiments.

### Odor delivery

For odor stimulus delivery for electrophysiological experiments, we used an eight-odor air dilution olfactometer. Approximately 1 sec prior to odor delivery a stream of Nitrogen was diverted through one of the odorant vials at a rate between 100 and 10 ml/min, and then merged into a clean air stream, flowing at a rate between 900 and 990 ml/min, thus providing 10 to 100 fold air dilution. Gas flows were controlled by mass flow controllers (Alicat MC series) with 0.5% accuracy. The odorized stream of 1000 ml/min was homogenized in a long thin capillary before reaching the final valve. Between stimuli, a steady stream of clean air with the same rate flowed to the odor port continuously, and the flow from the olfactometer was directed to an exhaust. During stimulus delivery, a final valve (four-way Teflon valve, Nresearch, SH360T042) switched the odor flow to the odor port, and diverted the clean airflow to the exhaust (Fig. S1). Temporal odor concentration profile was checked by mini PID (Aurora Scientific, model 200B). The concentration reached a steady state 95–210 ms (depending on a specific odor) after final valve opening. To minimize pressure shocks and provide temporally precise, reproducible, and fast odor delivery, we matched the flow impedances of the odor port and exhaust lines, and the flow rates from the olfactometer and clean air lines. As sniff activity was monitored in real time, the final valve was activated at the onset of exhalation, so that the odor reached steady-state concentration before the next inhalation. At the end of the odor delivery (duration 1-4 s) the final valve was deactivated, and Nitrogen flow was diverted from the odor vial to an empty line. Inter-odor delivery interval was 7-14 s, during which clean air was flowing through all Teflon tubing.

All odorants (see Table 1, purchased from Sigma-Aldrich) were diluted in mineral oil and stored in liquid phase in dark vials. The level of dilution of each odorant was estimated to achieve equal concentrations for all odorants of 0.075+0.01 μM after 10 fold air dilution^2,3,11^. Each vial contained 5 ml of mineral oil with diluted odorant and 45 ml of headspace.

For concentration series experiments for two odorants: 2HA and MEN, we changed dilution level in a range of approximately 2 orders of magnitude. The final desired concentrations were calibrated daily, immediately before the experiment began, and were achieved by tuning air dilution and matching PID signals between vials with different liquid dilutions.

Odor delivery system for imaging experiments was similar. However, due to differences in dilution procedure the matching concentrations for 2HA was 2x higher in imaging setup than that in electrophysiological and for MEN it was 1.8x lower.

### Olfactory bulb electrophysiology

MT cell spiking activity was recorded using 16-or 32-channel Si-probes (NeuroNexus, model: a2x2-tet-3mm-150-150-121(F16), Buzsaki32(F32)). Cells were recorded in the dorsal mitral cell layer. The identity of MT cells was established on the basis of criteria formulated in previous work^12^. The data were acquired using a 32-channel data acquisition system (HHMI Janelia Farm Research Campus, Applied Physics and Instruments Group,) with widely open broadband filters and sampling frequency of 19531 Hz.

### Recordings

#### Initial preparation

At the beginning of a recording session, a mouse was anesthetized with gas isofluorane and placed in the head-restraint setup. The running wheel was locked and a heating pad was placed under the animal. The lateral M72 glomerulus in either the right or left olfactory bulb was located using a fluorescent dissecting microscope, and the overlying bone was thinned. The open end of the fiber used for optical stimulation was positioned above the glomerulus, making contact with the thinned bone but without pressing on it.

A craniotomy was made just medial to the glomerulus, the dura removed and the silicon probe was inserted at an angle (25-45 degrees from vertical), driven by a digital micromanipulator (MP-285, Sutter Instruments). The insertion point was chosen so that at a depth of ∼300-500 μm from the brain surface, the tip of the most posterior shank in the probe would be roughly in line with the glomerulus in the medial/lateral axis. The anterior/posterior position was varied (following anatomical data from Liu & Urban, unpublished).

#### Search for MT cells putatively connected to M72 glomerulus

The anesthesia was removed and once the animal awakened, the probe was lowered to the EPL and advanced at ∼5μm intervals. At each position, a light pulse (0.5-15 mW power, 1-2 ms duration) was delivered to the glomerulus, triggered on the onset of inhalation. The peri-stimulus activity on all sites of the Si-probe was monitored. A spiking increase with short latency after the light pulse (below 20 ms, typically 5-10 ms) indicated the presence of a cell receiving input from the stimulated glomerulus. If no light responsive cells were found upon reaching a depth of ∼700-800 μm, the electrode was raised, reinserted, and the search repeated.

#### Recording odor responses

After locating a putative M72 MT cell, odor recording session was initiated. Multiple odorant stimuli with fixed concentrations or two odorant stimuli with multiple concentrations were presented pseudo-randomly with 7-14 s inter-trial interval. After every 2-4 odor trials, a light pulse was delivered, to re-confirm the presence of the M72 MT cell. For each odor stimulus, 20-35 trials were collected.

All sites of the Si probe were used to monitor activity of other, non light-responsive units, during M72 MT recording sessions. In addition, to increase the pool of non-M72 MT cell (other cells), we anesthetized the animal again, performed a new craniotomy, and placed the probe at a new site, usually further anterior, and performed recordings with the same stimulus set.

### Spike extraction

Acquired electrophysiological data were filtered and spike sorted. We used the Klusta suite software package for spike detection and spike sorting^9,13^ and software written by E.M.A. and D.R.

### Identification of M72-MT cells

We defined MT cells functionally connected to the M72 glomerulus as units that displayed an excitatory, short latency response to light stimulation (1 ms, 5-15 mW) of the ChR2-expressing M72 glomerulus. While in general it is difficult to establish monosynaptic connectivity using optogenetic stimulation^14^, we capitalized on the known anatomy of the olfactory bulb: MT cells receive excitatory input from a single glomerulus, and interactions between MT cells connected to different glomeruli are inhibitory^15,16^.

We compared the PSTHs of MT cells with and without light stimulation. PSTHs with 4 ms temporal bins were referenced to the onset of inhalation at the onset of inhalation, when the light stimulation was presented. The MT cell was considered light responsive if light-evoked activity exceeded activity in the no-light condition by at least one standard deviation, in at least one 4 ms temporal bin, within 50 ms after the onset of the light pulse. The latency was estimated as the first time point when such a deviation occurred. The distribution of latencies is shown in Fig. 1e. The majority of the responses (1.5 IQR) occurred with latencies shorter than 22 ms. Two cells responded with latencies larger than 1.5 IQR, 22 ms, and were were removed from the pool of cells used in this study.

### Snifflet analysis of the response profiles

We build probabilistic models to describe the encoding of odor stimuli by MT neurons. These models take the form of Generalized Linear Models (GLMs)^17,18^, and describe a generative model for the spiking data. For a given cell and a given odor (or the baseline condition, with no odor presentation), we assume that the firing rate of the cell changes over the course of the sniff according to a specific temporal pattern, which we call a “snifflet”. Given that the observed spiking patterns during individual sniffs depend on the inhalation duration (Fig. 2d), we build this dependency into the model. In particular, we assume that during an individual sniff, the firing rate is generated by temporally dilating the snifflet by a sniff-dependent factor. The value of this dilation factor, α, depends on the duration of the inhalation phase of that sniff. More formally, we write the rate *r(t)* as

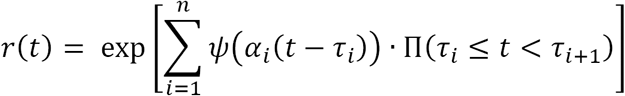

where *Ψ*(*t*) is the odor-evoked snifflet, *α*_*i*_ is the temporal dilation factor for the *i*^*th*^ sniff, *τ*_*i*_ is the onset time of the *i*^*th*^ sniff, *n* is the total number of sniffs and Π(·) is an indicator function, such that the response pattern is reset at the onset of the next sniff. Removing the indicator function, so that snifflets from successive odors overlap, does not qualitatively change any of the major results in the main text.

The model requires a choice of dilation factors, *α*_*i*_, for each sniff. Motivated by the work of *Shusterman et. al.*^9^, we fix the dilation factors as the reciprocal of the inhalation durations, 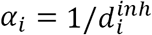, where 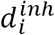 is a duration of the inhalation phase of the sniff cycle. This produced better model fits than alternative choices, such as dilating with the reciprocal of the full sniff durations, separate dilation for the inhalation and rest-of-sniff components of the response, or fixing *α*_*i*_ = 1 (i.e. no temporal dilation) throughout (Fig. S4).

The free parameters of the model are the snifflet time course, *Ψ*(*t*), for each cell and odor. We parameterize the snifflets as a length-KD vector (for integers K and D), such that the first D components represent the evolution of the cell’s firing rate during the inhalation period, and the remaining (K-1)D components represent the evolution of the cell’s firing rate during the remainder of the sniff. The integer D thus defines the sampling resolution for the snifflet, and K the relative duration of post-inhalation response to model. We use D=30 and K=4 in the main text, but other values produced similar results.

We also placed priors over the components of *Ψ*(*t*), to constrain the snifflets would evolve smoothly in time. We used the Automatic Smoothness Determination prior^19^ and learned the hyper-parameters via evidence optimization^20,21^. Including the prior dramatically increased the quality of model predictions on held-out data (Fig. S2).

We solve for *Ψ*(*t*), by maximizing its posterior probability. Given a point estimate of the hyper-parameters, and using a fixed scheme for determining *α* (above), this is a convex problem^22^, which we solve using conventional Newton methods. We approximate the posterior on *Ψ*(*t*) using a Laplace approximation. For the purposes of illustration, we show the snifflets in the main text in their exponentiated form (i.e., in terms of firing rate, rather than log firing rate). Where error bars on individual snifflets are shown (Fig. 2e), the shaded areas illustrate only the marginal variance of the approximate posterior at each time point, rather than the joint covariance across time. Statistical comparisons between odor-evoked and baseline snifflets (Figs. 4-5) were performed in log firing rate space; we again consider only the marginal variance at each time point.

The snifflet model provides a parameterization of how inhalation duration (a nuisance variable) affects spiking response, allowing us to factor this relationship out from our results and study the differences across odors and cells. To verify that our results did not depend on the particulars of the snifflet model, we fitted the spiking data without adjusting for variations in sniff duration Fig. S4).

### Data availability

The datasets generated during and/or analysed during the current study are available from the corresponding author on reasonable request.

### Code availability

Code developed for this work is available in the following github repositories: https://github.com/admiracle/Voyeur (stimulus delivery system control); https://github.com/zekearneodo/ephys-tools (post-recording data preparation, pre-processing and initial sniff analysis); https://github.com/rabbitmcrabbit/snifflet (snifflet analysis).

---

**Figure S1.**
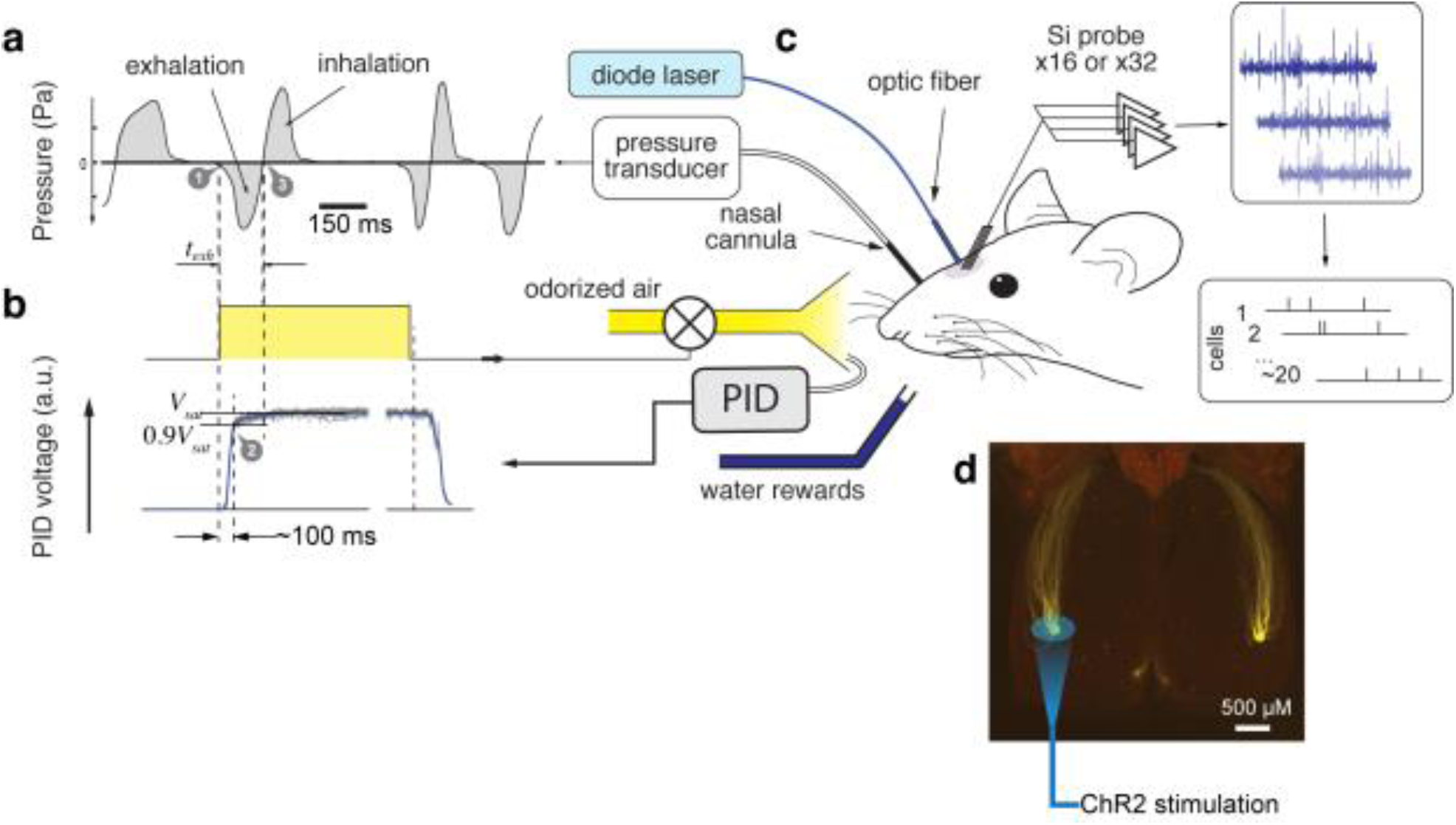
Experimental setup. The mouse is implanted with a sniffing cannula for pressure measurement and a head bar for head fixation (not shown). After recovery, the mouse is acclimated to the experimental apparatus. **(c)** At the beginning of the electrophysiological recording session, the animal is anesthetized with isofluorane. The YFP-expressing M72 glomerulus is located and the optical fiber placed above. A small craniotomy is made next to the glomerulus and a 16 or 32 site Si-probe is inserted, targeting the mitral cell layer near the glomerulus. The electrode is connected to the data acquisition system via a 32 channel headstage. Raw data is stored in the computer and spikes are sorted offline. All electrophysiological recordings are done after full recovery from anesthesia. **(a)** A typical pressure waveform from the nasal cavity is shown. **(b)** During the experimental session, a final valve, which delivers odors, is triggered at the onset of exhalation (**a**, time point 1). Such triggering ensures that when a mouse starts inhaling an odor (time point 3), the odor concentration has reached a steady state (time point 2). The odor delivery system is calibrated prior to every experimental session using photoionization detector (PID). The typical profiles of PID traces for repeatable odor delivery with a constant concentration are shown at (**b**): thin gray lines are individual trials, and blue line is an average profile. During the first 100 ms from the final valve trigger, odor concentration reaches 90% of its asymptotic value. For a majority of trials, exhalation duration *t*_*exh*_ is longer than 100 ms. (**d**) Dorsal view of olfactory bulbs of M72-ChR2-YFP mouse. Axons of M72 receptor neurons, co-expressing ChR2 and YFP converge to dorsal glomeruli in the left and right bulbs. Optical fiber connected to blue laser is positioned above dorsal M72 glomerulus.

**Figure S2.**
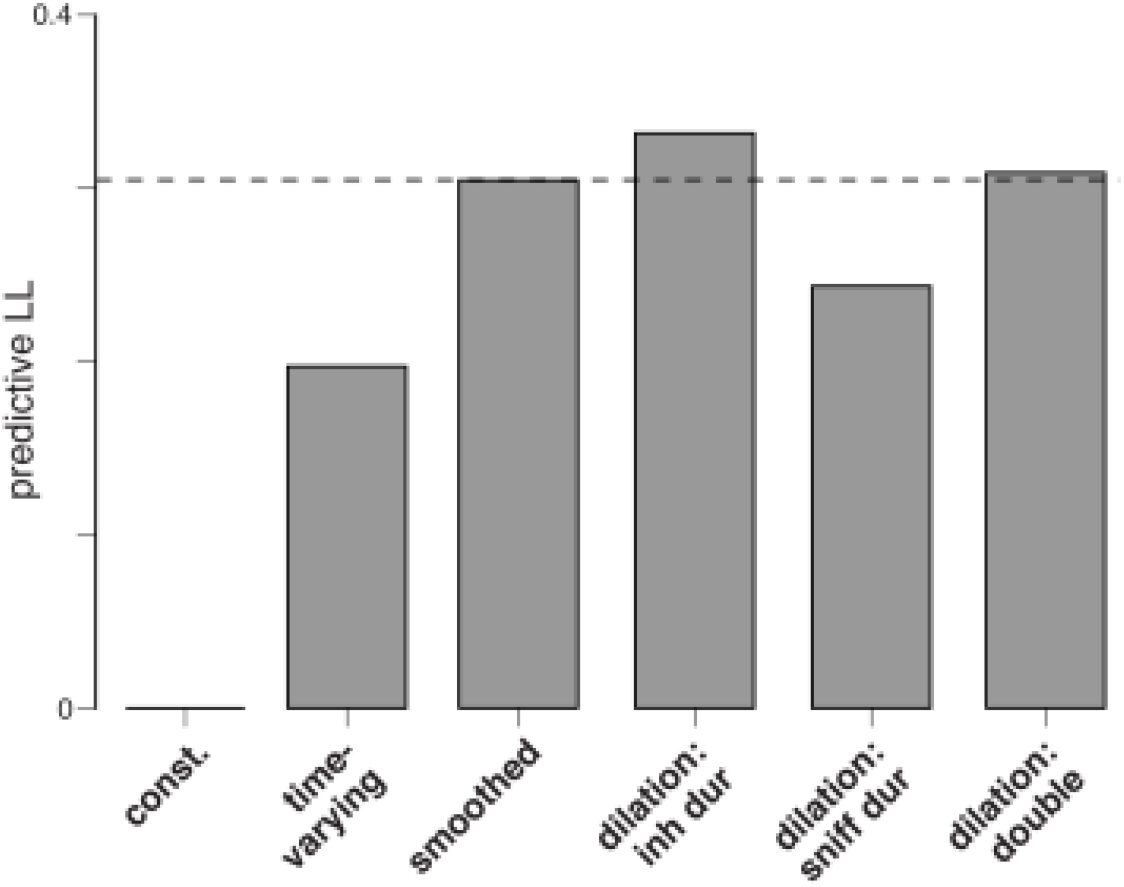
Quality of model fits, measured as log likelihood (LL) on held-out data. In the main text, we describe the fitting of snifflet models to odor-evoked responses of MT cells. Here we quantify the quality of these fits, and justify the modelling decisions made. Bars show the predictive LL (per second) on held-out data of the fitted models, averaged across MT cells and odors. Note that these scores are cross-validated: improvements indicate that the models are explaining structure in the data, rather than overfitting. Values are expressed relative to the constant model (left), i.e. where the firing rate of each neuron is assumed to be constant over the duration of the sniff. The other bars show four snifflet models. (**time-varying)**: odor responses described via an unsmoothed PSTH with a ridge prior, and no temporal dilation. This shows the performance of a naive, sniff-aligned model of odor-evoked responses. (**smoothed)**: Same, but with a smoothed PSTH, with a learned smoothing time constant for each cell/odor. (**dilation: inh dur**): the smoothed snifflet model, where the snifflet is also temporally dilated for each sniff based on inhalation duration. This is the model used in the main text. (**dilation: sniff dur**): the smoothed snifflet model, where the snifflet is temporally dilated for each sniff based on the total sniff duration. This has poorer predictive performance than the model in the main text. (**dilation: double**): the smoothed snifflet model, where the snifflet is temporally diluted at two intervals: sniff duration and the rest of the sniff. This model performed worse than pure inhalation duration dilation model.

**Figure S3.**
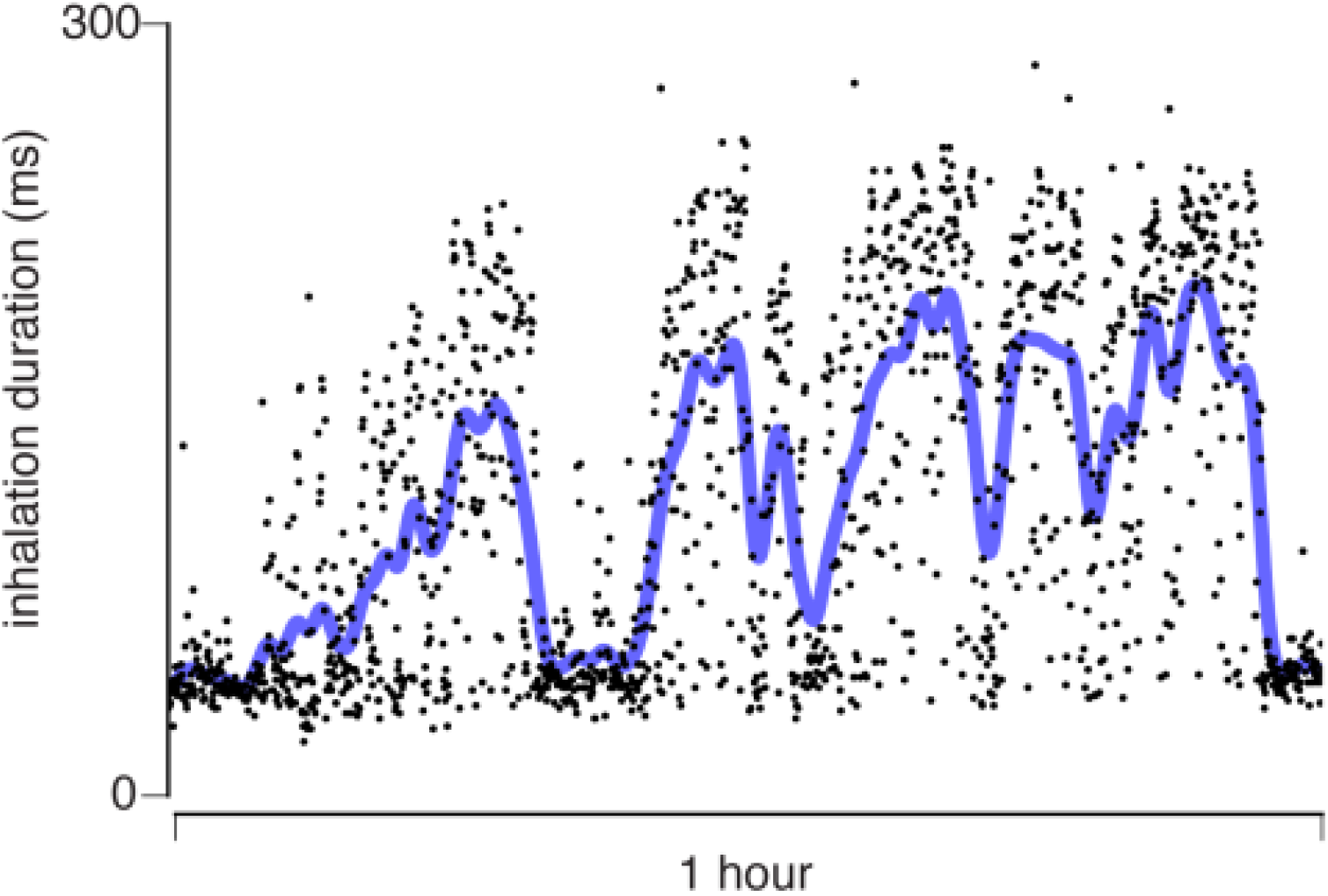
Patterns of sniffing in an individual animal over an hour. Black dots show the inhalation duration of successive sniffs during one recording session; thick blue line shows a temporally smoothed version of the signal. The rarer fast sniffs predominantly occur during isolated periods, here seen at the beginning and end of the session, and at about 20 minutes in.

**Figure S4.**
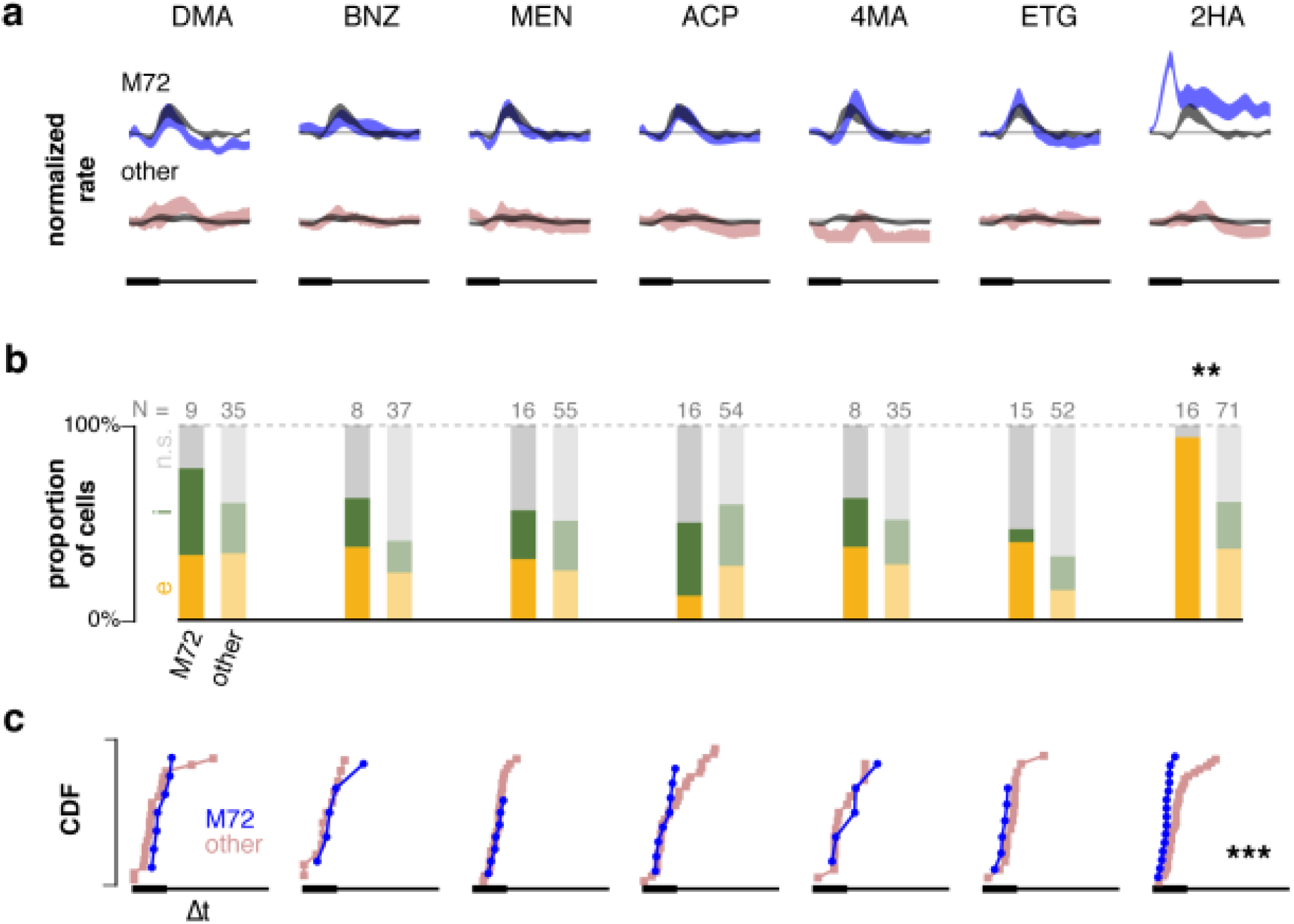
Results shown in Figure 4, obtained with the “no dilation” snifflet model (Figure S2, third column). The results are qualitatively identical to those shown in Figure 4. Similar results are obtained for the results in Figure 5 with this model as well (not shown).

**Figure S5.**
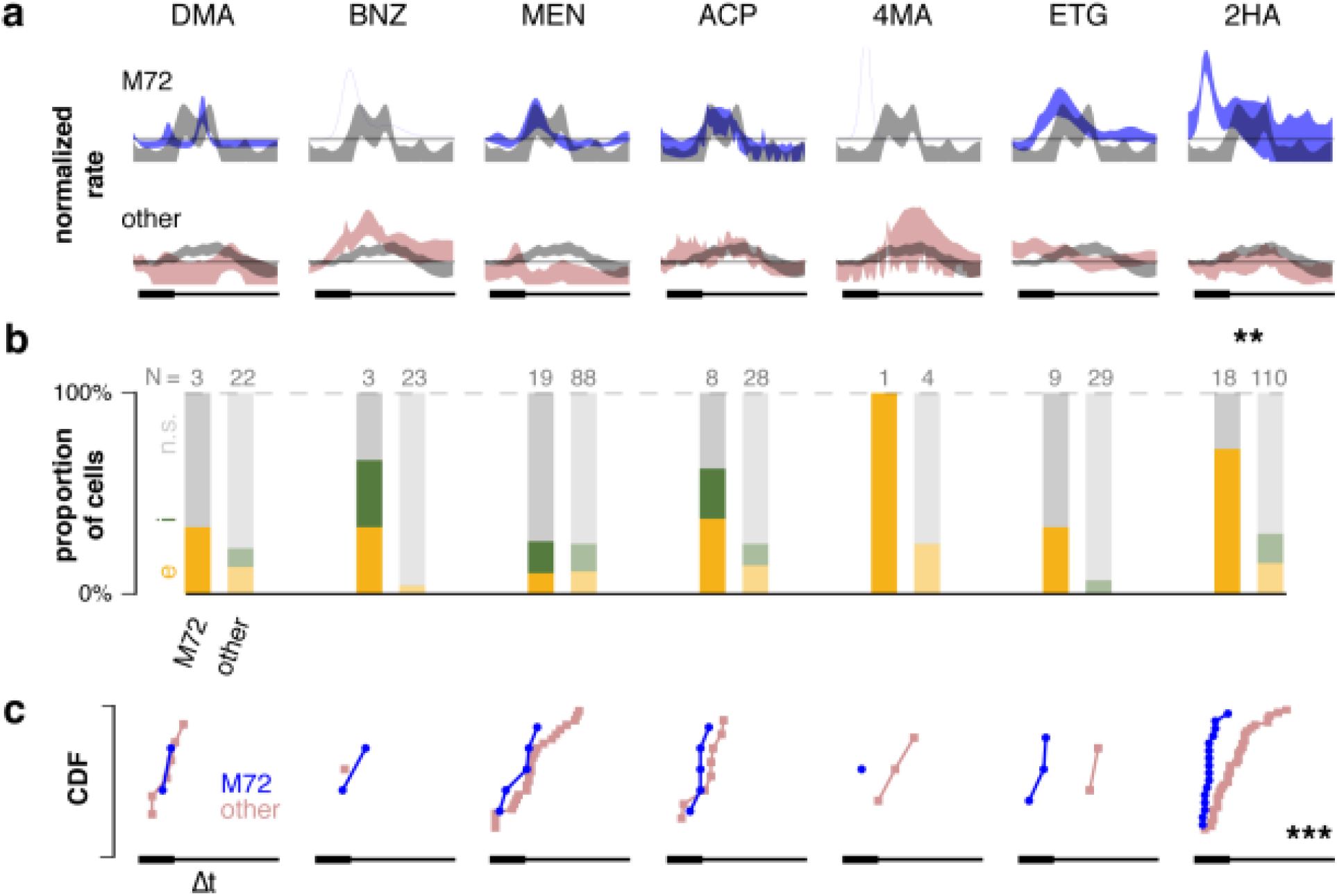
Results shown in Figure 4, when considering only fast sniffs (inhalation durations shorter than 100 ms) rather than slow sniffs. As explained in the main text, fast sniffs were rarer (<25%), and seemed to occur as part of a distinct behavioral state of the animals (Figure S2). We therefore focused exclusively on slower sniffs in the main text. Here we repeat the analysis of Figure 4 on the fast sniffs only. As fast sniffs were rare for many animals, there was not enough data to analyse for most cells. However, the same trends were evidence as in Figure 4. The yield for the concentration experiments in Figure 5 was even smaller, making a comparable analysis of responses to these stimuli during fast sniffs impractical.

**Figure S6.**
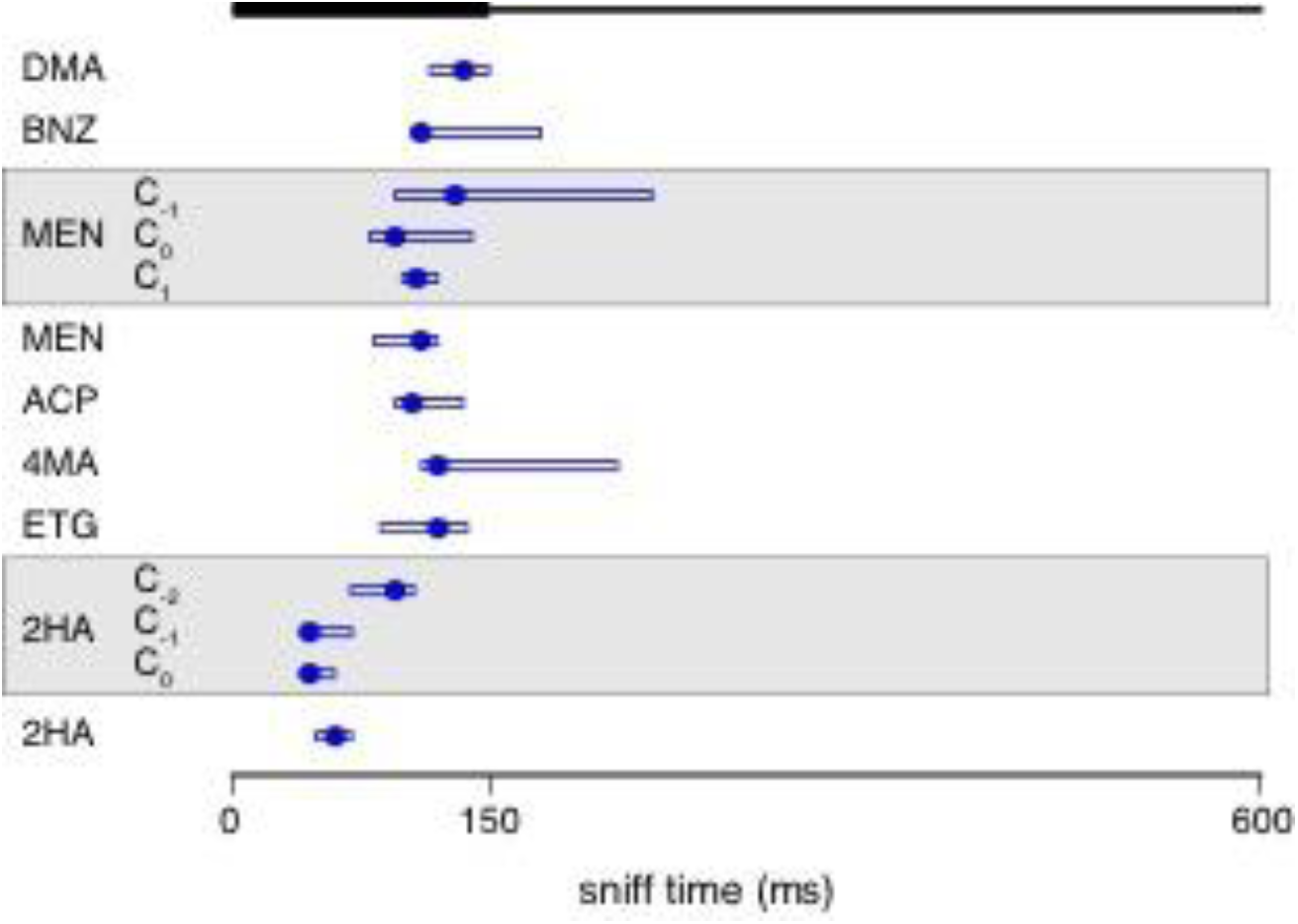
Average latency of first significant response to each odor for M72-MT cells. Dots show the median first-response latency across cells; bars show the interquartile range. The odors against the white background summarize the latency distributions shown in Figure 4c; the odors against the grey background summarize the latency distributions shown in Figure 5c.

**Figure S7.**
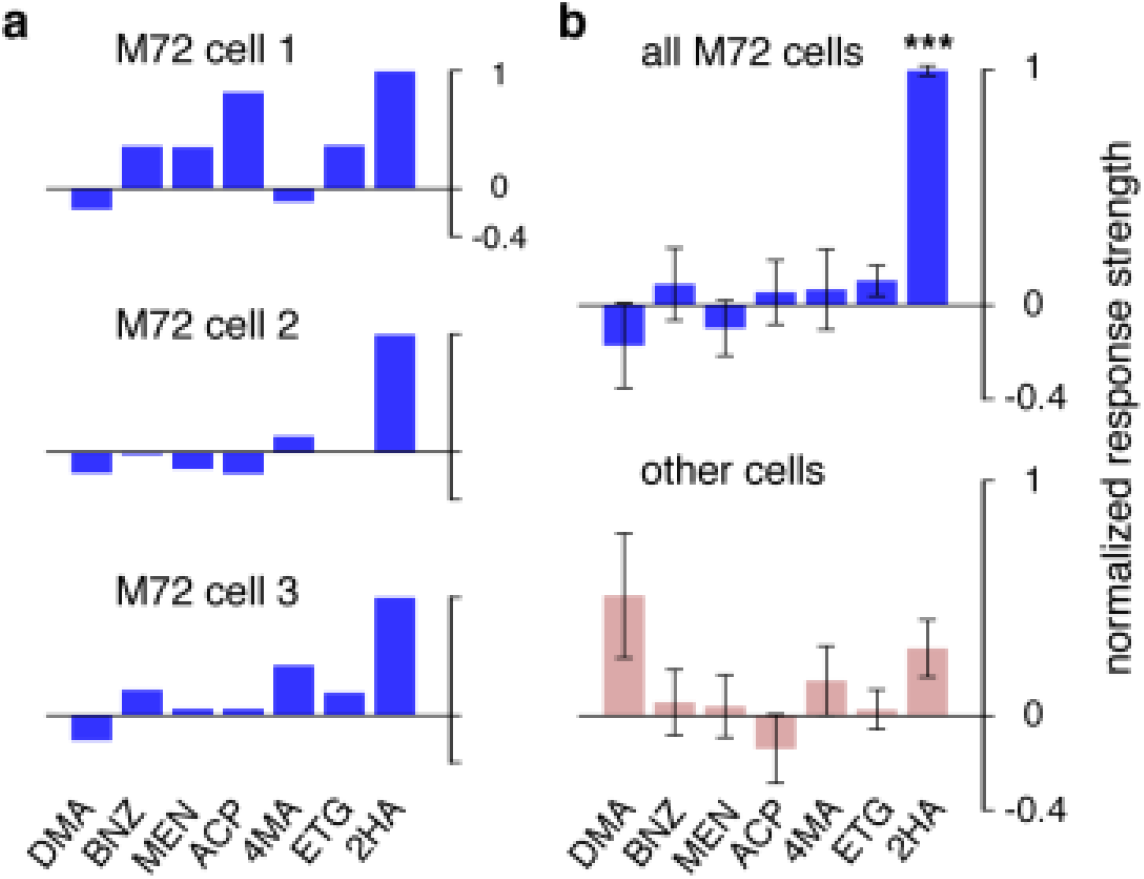
Stereotypy in responses to strongest ligand is also observed in odor-evoked rate changes. **(a)** Odor response profiles, calculated by mean change in spike count over the duration of the entire sniff, shown for three M72-MT cells to the 7 odors. Responses are normalized such that 0 (horizontal line) is the spike count during baseline (air), and 1 is the maximal spike count across odors. **(b)** Odor response profiles, averaged across cells; for the M72-MT population (top) and the generic MT population (bottom), showing the M72-MT response is significantly higher than the generic MT response only when the stimulus is 2HA (Wilcoxon signed-rank test). Error bars = S.E.M.

## References

1. Mombaerts, P. et al. Visualizing an olfactory sensory map. Cell 87, 675–686 (1996).

2. Bozza, T., Feinstein, P., Zheng, C. & Mombaerts, P. Odorant receptor expression defines functional units in the mouse olfactory system. J Neuroscience 22, 3033–3043 (2002).

3. Shepherd, G. M. The Synaptic Organization of the Brain. (Oxford University Press, USA, 2004).

4. Spors, H. & Grinvald, A. Spatio-temporal dynamics of odor representations in the mammalian olfactory bulb. Neuron 34, 301–315 (2002).

5. Carey, R. M., Verhagen, J. V., Wesson, D. W., Pírez, N. & Wachowiak, M. Temporal structure of receptor neuron input to the olfactory bulb imaged in behaving rats. J Neurophysiol 101, 1073–1088 (2009).

6. Shusterman, R., Smear, M. C., Koulakov, A. A. & Rinberg, D. Precise olfactory responses tile the sniff cycle. Nat Neurosci 14, 1039–1044 (2011).

7. Cury, K. M. & Uchida, N. Robust Odor Coding via Inhalation-Coupled Transient Activity in the Mammalian Olfactory Bulb. Neuron 68, 570–585 (2010).

8. Tan, J., Savigner, A., Ma, M. & Luo, M. Odor information processing by the olfactory bulb analyzed in gene-targeted mice. Neuron 65, 912–926 (2010).

9. Kikuta, S., Fletcher, M. L., Homma, R., Yamasoba, T. & Nagayama, S. Odorant response properties of individual neurons in an olfactory glomerular module. Neuron 77, 1122–1135 (2013).

10. Dhawale, A. K., Hagiwara, A., Bhalla, U. S., Murthy, V. N. & Albeanu, D. F. Non-redundant odor coding by sister mitral cells revealed by light addressable glomeruli in the mouse. Nat Neurosci 13, 1404–1412 (2010).

11. Zhang, J., Huang, G., Dewan, A., Feinstein, P. & Bozza, T. Uncoupling stimulus specificity and glomerular position in the mouse olfactory system. Mol. Cell. Neurosci. 51, 79–88 (2012).

12. Potter, S. M. et al. Structure and emergence of specific olfactory glomeruli in the mouse. J Neuroscience 21, 9713–9723 (2001).

13. Smear, M., Resulaj, A., Zhang, J., Bozza, T. & Rinberg, D. Multiple perceptible signals from a single olfactory glomerulus. Nat Neurosci 16, 1687–1691 (2013).

14. Buonviso, N., Amat, C. & Litaudon, P. Respiratory modulation of olfactory neurons in the rodent brain. Chem Senses 31, 145–154 (2006).

15. Paninski, L. Maximum likelihood estimation of cascade point-process neural encoding models.Network 15, 243–262 (2004).

16. Park, M. & Pillow, J. W. Receptive field inference with localized priors. PLoS Comput Biol (2011).

17. Sirotin, Y. B., Shusterman, R. & Rinberg, D. Neural Coding of Perceived Odor Intensity. eNeuro 2,(2015).

18. Padmanabhan, K. & Urban, N. N. Intrinsic biophysical diversity decorrelates neuronal firing while increasing information content. Nat Neurosci 13, 1276–1282 (2010).

19. Angelo, K. et al. A biophysical signature of network affiliation and sensory processing in mitral cells. Nature 488, 375–378 (2012).

20. Fukunaga, I., Berning, M., Kollo, M., Schmaltz, A. & Schaefer, A. T. Two distinct channels of olfactory bulb output. Neuron 75, 320–329 (2012).

21. Egger, V. & Urban, N. N. Dynamic connectivity in the mitral cell-granule cell microcircuit. Semin Cell Dev Biol 17, 424–432 (2006).

22. Kato, H. K., Gillet, S. N., Peters, A. J., Isaacson, J. S. & Komiyama, T. Parvalbumin-expressing interneurons linearly control olfactory bulb output. Neuron 80, 1218–1231 (2013).

23. Chen, T.-W., Lin, B.-J. & Schild, D. Odor coding by modules of coherent mitral/tufted cells in the vertebrate olfactory bulb. Proc Natl Acad Sci USA 106, 2401–2406 (2009).

24. Whitesell, J. D., Sorensen, K. A., Jarvie, B. C., Hentges, S. T. & Schoppa, N. E. Interglomerular lateral inhibition targeted on external tufted cells in the olfactory bulb. J Neuroscience 33, 1552–1563 (2013).

25. Fantana, A. L., Soucy, E. R. & Meister, M. Rat olfactory bulb mitral cells receive sparse glomerular inputs. Neuron 59, 802–814 (2008).

26. Margrie, T. W. & Schaefer, A. T. Theta oscillation coupled spike latencies yield computational vigour in a mammalian sensory system. J Physiol (Lond) 546, 363–374 (2003).

27. Cang, J. & Isaacson, J. S. In vivo whole-cell recording of odor-evoked synaptic transmission in the rat olfactory bulb. J Neuroscience 23, 4108–4116 (2003).

28. Mozell, M. M. Evidence for a chromatographic model of olfaction. J Gen Physiol 56, 46–63 (1970).

29. Jiang, J. & Zhao, K. Airflow and nanoparticle deposition in rat nose under various breathing and sniffing conditions: a computational evaluation of the unsteady effect. J Aerosol Sci 41, 1030–1043 (2010).

30. Ghatpande, A. S. & Reisert, J. Olfactory receptor neuron responses coding for rapid odor sampling. J Physiol (Lond) (2011). doi:10.1113/jphysiol.2010.203687.

31. Chen, W. R. & Shepherd, G. M. The olfactory glomerulus: a cortical module with specific functions. J Neurocytol 34, 353–360 (2005).

32. Jeanne, J. M. & Wilson, R. I. Convergence, Divergence, and Reconvergence in a Feedforward Network Improves Neural Speed and Accuracy. Neuron (2015). doi:10.1016/j.neuron.2015.10.018.

33. van Drongelen, W., Holley, A. & Døving, K. B. Convergence in the olfactory system: quantitative aspects of odour sensitivity. J Theor Biol 71, 39–48 (1978).

34. Luo, M. & Katz, L. C. Response correlation maps of neurons in the mammalian olfactory bulb. Neuron 32, 1165–1179 (2001).

35. Willhite, D. C. et al. Viral tracing identifies distributed columnar organization in the olfactory bulb. Proc Natl Acad Sci USA 103, 12592–12597 (2006).

36. Vucinic, D., Cohen, L. B. & Kosmidis, E. K. Interglomerular center-surround inhibition shapes odorant-evoked input to the mouse olfactory bulb in vivo. J Neurophysiol 95, 1881–1887 (2006).

37. Urban, N. N. Lateral inhibition in the olfactory bulb and in olfaction. Physiol Behav 77, 607–612 (2002).

38. Kuffler, S. W. Discharge patterns and functional organization of mammalian retina. J Neurophysiol 16, 37–68 (1953).

39. Hartline, H. K. & Ratliff, F. Inhibitory interaction of receptor units in the eye of Limulus. J Gen Physiol 40, 357–376 (1957).

40. Knudsen, E. I. & Konishi, M. Center-surround organization of auditory receptive fields in the owl. Science 202, 778–780 (1978).

41. Sutter, M. L., Schreiner, C. E., McLean, M., O’connor, K. N. & Loftus, W. C. Organization of inhibitory frequency receptive fields in cat primary auditory cortex. J Neurophysiol 82, 2358–2371 (1999).

42. Hellweg, F. C., Schultz, W. & Creutzfeldt, O. D. Extracellular and intracellular recordings from cat’s cortical whisker projection area: thalamocortical response transformation. J Neurophysiol 40, 463–479 (1977).

43. Laskin, S. E. & Spencer, W. A. Cutaneous masking. II. Geometry of excitatory andinhibitory receptive fields of single units in somatosensory cortex of the cat. J Neurophysiol 42, 1061–1082 (1979).

44. Sur, M. Receptive fields of neurons in areas 3b and 1 of somatosensory cortex in monkeys. Brain Res 198, 465–471 (1980).

45. Cleland, T. A. & Sethupathy, P. Non-topographical contrast enhancement in the olfactory bulb. BMC Neurosci 7, 7 (2006).

46. Olsen, S. R., Bhandawat, V. & Wilson, R. I. Divisive normalization in olfactory population codes. Neuron 66, 287–299 (2010).3

## References

1. Tian, L. et al. Imaging neural activity in worms, flies and mice with improved GCaMP calcium indicators. Nat Methods 6, 875–881 (2009).

2. Smear, M., Resulaj, A., Zhang, J., Bozza, T. & Rinberg, D. Multiple perceptible signals from a single olfactory glomerulus. Nat Neurosci 16, 1687–1691 (2013).

3. Zhang, J., Huang, G., Dewan, A., Feinstein, P. & Bozza, T. Uncoupling stimulus specificity and glomerular position in the mouse olfactory system. Mol. Cell. Neurosci. 51, 79–88 (2012).

4. Bozza, T., McGann, J. P., Mombaerts, P. & Wachowiak, M. In vivo imaging of neuronal activity by targeted expression of a genetically encoded probe in the mouse. Neuron 42, 9–21 (2004).

5. Bunting, M., Bernstein, K. E., Greer, J. M., Capecchi, M. R. & Thomas, K. R. Targeting genes for selfexcision in the germ line. Genes Dev. 13, 1524–1528 (1999).

6. Shaner, N. C. et al. Improved monomeric red, orange and yellow fluorescent proteins derived from Discosoma sp. red fluorescent protein. Nat. Biotechnol. 22, 1567–1572 (2004).

7. Potter, S. M. et al. Structure and emergence of specific olfactory glomeruli in the mouse. J Neuroscience 21, 9713–9723 (2001).

8. Osborne, J. E. & Dudman, J. T. RIVETS: A Mechanical System for In Vivo and In Vitro Electrophysiology and Imaging. PLoS ONE 9, e89007 (2014).

9. Shusterman, R., Smear, M. C., Koulakov, A. A. & Rinberg, D. Precise olfactory responses tile the sniff cycle. Nat Neurosci 14, 1039–1044 (2011).

10. Sirotin, Y. B., Shusterman, R. & Rinberg, D. Neural Coding of Perceived Odor Intensity. eNeuro 2, (2015).

11. Wojcik, P. T. & Sirotin, Y. B. Single scale for odor intensity in rat olfaction. Curr Biol 24, 568–573 (2014).

12. Rinberg, D., Koulakov, A. & Gelperin, A. Sparse odor coding in awake behaving mice. J Neuroscience 26, 8857–8865 (2006).

13. Hazan, L., Zugaro, M. & Buzsáki, G. Klusters, NeuroScope, NDManager: a free software suite for neurophysiological data processing and visualization. J Neurosci Methods 155, 207–216 (2006).

14. Lima, S. Q., Hromádka, T., Znamenskiy, P. & Zador, A. M. PINP: a new method of tagging neuronal populations for identification during in vivo electrophysiological recording. PLoS ONE 4, e6099 (2009).

15. Urban, N. N. & Sakmann, B. Reciprocal intraglomerular excitation and intra-and interglomerular lateral inhibition between mouse olfactory bulb mitral cells. J Physiol (Lond) 542, 355–367 (2002).

16. Shepherd, G. M. The Synaptic Organization of the Brain. (Oxford University Press, USA, 2004).

17. Truccolo, W., Eden, U. T., Fellows, M. R., Donoghue, J. P. & Brown, E. N. A point process framework for relating neural spiking activity to spiking history, neural ensemble, and extrinsic covariate effects. J Neurophysiol 93, 1074–1089 (2005).

18. Pillow, J. W. et al. Spatio-temporal correlations and visual signalling in a complete neuronal population. Nature 454, 995–999 (2008).

19. Linden, M. Evidence optimization techniques for estimating stimulus-response functions. Neural Information Processing Systems 15, 317 (2003).

20. Park, M. & Pillow, J. W. Receptive field inference with localized priors. PLoS Comput Biol (2011).

21. Park, M. & Pillow, J. W. Bayesian inference for low rank spatiotemporal neural receptive fields. 2688– 2696 (2013).

22. Paninski, L. Maximum likelihood estimation of cascade point-process neural encoding models. Network 15, 243–262 (2004).

